# Influence of Disease-Causing Mutations and Ivacaftor on the Dynamics of the Cystic Fibrosis Transmembrane Conductance Regulator (CFTR) Protein

**DOI:** 10.1101/2025.07.07.663434

**Authors:** Diana-F. Veselu, Deborah K. Shoemark, Marc W. van der Kamp

**Affiliations:** School of Biochemistry, University of Bristol, Bristol BS8 1TD, UK

## Abstract

Cystic fibrosis (CF), one of the most common life-shortening autosomal recessive diseases, arises from mutations in the cystic fibrosis transmembrane conductance regulator (CFTR) gene and leads to severe respiratory and digestive dysfunction. CFTR-targeting therapies such as ivacaftor have transformed CF care by directly modulating CFTR function. However, the precise mechanism by which ivacaftor alters CFTR’s conformational dynamics remains incompletely understood. In this study, we refined and employed a CFTR model derived from the phosphorylated, ATP- and ivacaftor-bound cryo-EM structure (PDB ID 6O2P). Through 32.5 μs of molecular dynamics simulations, we investigated the structural and dynamic effects of the common F508del mutation and the gating mutations G551D, G1349D and S549N. Our results show that mutations within the ATP-binding regions cause local rearrangements at the nucleotide-binding sites, affecting the interface between the nucleotide-binding domains (NBDs) relative to wild-type CFTR, rather than abolishing ATP binding entirely. We further indicate that, for the NBDs to approach one another productively, protonation of the aspartate side chains in G551D and G1349D CFTR is necessary to mitigate charge repulsion. The F508del mutation induces increased rigidity within the protein core, impeding the rearrangement of transmembrane helices required for channel opening. Ivacaftor does not directly perturb the ATP-binding sites, consistent with emerging evidence that it exerts its effects allosterically on regions distal to the nucleotide-binding sites. Collectively, our findings enhance our understanding of CFTR dynamics and provide a versatile framework for exploring the molecular effects of disease-causing mutations and evaluating potential therapeutic agents.

## Introduction

Cystic Fibrosis (CF) is a life-shortening autosomal recessive disorder that affects multiple organ systems, leading to significant morbidity and reduced life expectancy.^1^ A hallmark of CF is the accumulation of thick, sticky mucus that obstructs epithelia-lined ducts and tubes, particularly in the lungs and pancreas, resulting in chronic infections, progressive lung damage, and severe digestive complications from pancreatic-duct obstruction.^2^

CF is caused by mutations in the cystic fibrosis transmembrane conductance regulator (CFTR) gene, which encodes a chloride/bicarbonate channel in epithelial membranes that regulates fluid and electrolyte balance.^2^ CFTR belongs to the ATP-binding cassette (ABC) protein family, but unlike typical ABC proteins, CFTR functions as an ATP-gated ion channel.^3^ Its architecture comprises two membrane-spanning domains (MSD1 and MSD2), two nucleotide-binding domains (NBD1 and NBD2), and a distinct, intrinsically disordered regulatory (R) domain. In the dephosphorylated state, the R domain obstructs channel activation by preventing the close association or dimerisation of the NBDs, a process we refer to as dimerization for simplicity. Once phosphorylated by cAMP-dependent protein kinases, this inhibition is lifted, enabling ATP binding and subsequent NBD dimerisation that drives channel gating.^4^

To date, over 2,000 CFTR mutations have been identified,^5^ each disrupting CFTR activity through one or more molecular mechanisms, including impaired protein trafficking, defective channel gating, reduced membrane stability, or aberrant protein synthesis.^6^ The most common mutation, F508del (a deletion of phenylalanine at position 508), exemplifies this pleiotropy, causing protein misfolding, gating defects, and decreased cell-surface stability.^6^

The complexity and variability of CFTR mutations present challenges for successful CF treatment. Effective therapies should not only relieve symptoms, but also correct the specific molecular defects associated with each mutation. Achieving this goal likely requires detailed understanding of CFTR’s structure, dynamical behaviour, interactions with modulators, and the influence of its membrane environment. In recent years, CFTR modulators (small molecules that directly restore CFTR function) have transformed the CF therapeutic landscape.^7^ These include potentiators (which enhance channel gating), correctors (which rescue protein folding/trafficking), stabilisers (which prolong membrane retention), read-through agents (which bypass premature stop codons), and amplifiers (which increase overall CFTR protein production).^7^ Combination regimens of one or two correctors with a potentiator have shown substantial clinical benefit for patients with responsive genotypes.^7^

High-resolution cryo-electron microscopy (cryo-EM) studies have provided detailed static snapshots of CFTR and its drug-bound conformations. The first high-resolution cryo-EM structure of zebrafish CFTR in its dephosphorylated, ATP-free state,^8^ laid the foundation for subsequent human CFTR structures in various functional states, including ATP-bound and drug-bound forms.^9–12^ Despite these advances, our understanding of CFTR’s conformational dynamics remains limited. In particular, how modulators such as ivacaftor affect CFTR structure and function, especially in mutant backgrounds, and the role of the surrounding lipid membrane in stabilising CFTR and facilitating drug binding, are not yet fully resolved.

In this study, we use molecular dynamics simulations to investigate the behaviour of membrane-embedded CFTR in its phosphorylated, ATP-bound state. We constructed a detailed atomistic model of human CFTR based on the cryo-EM structure of the phosphorylated, ATP- and ivacaftor-bound state (PDB ID 6O2P)^9^ and performed extensive simulations on wild-type (WT) CFTR, the common F508del mutant, and the gating mutations G551D, G1349D, and S549N. For G551D and G1349D, both protonated (G551DH, G1349DH) and unprotonated aspartate (G551D, G1349D) forms were simulated. Each system was simulated under both ivacaftor-free and ivacaftor-bound conditions to examine how these mutations and ivacaftor influence CFTR structure and dynamics. Such detailed molecular dynamics simulations of membrane proteins can provide insights at the atomic level that are difficult to reveal from experimental methods alone.^13^

Our simulations extend structural data to reveal mechanistic insights into CFTR’s dynamic behaviour. We find that the glycine-to-aspartate mutations G551D and G1349D, which lie adjacent to the ATP-binding sites, likely require aspartate protonation to facilitate stable NBD dimerisation required for activity. Although these mutations, along with S549N, do not fully abolish ATP binding, they impair channel gating through allosteric disruption beyond the nucleotide-binding pockets. The F508del mutation increases core rigidity, hindering the conformational transitions needed for channel opening. Finally, our results support the growing view that ivacaftor enhances gating primarily via allosteric modulation of the membrane-spanning domains rather than by directly affecting the ATP-binding sites. These findings contribute to a deeper, mutation-specific understanding of CFTR dynamics and may inform future therapeutic strategies.

## Methods

### System preparation

We constructed a human CFTR model (Figure 1), based on the cryo-EM structure of the phosphorylated, ATP- and ivacaftor-bound state (PDB ID 6O2P).^9^ Missing loops (residues 410-436, 890-899, 1174-1201) were grafted from the AlphaFold2 model AF-P13569-F1-v4 (Figure S1).^14, 15^ A cryo-EM structure was chosen as this provides the only full-length experimental structure of CFTR while X-ray crystallography has resolved only isolated domains, such as the NBD1 and truncated NBD1 constructs.^16^ The regulatory (R) domain (>200 residues) was omitted, as it is almost certainly intrinsically disordered: there is no electron density for the majority of this domain in cryo-EM structures, and the AlphaFold2 model shows very low prediction confidence (pLDDT < 50 and PAE > 25 for the majority of the domain). Its expected wide conformational ensemble would thus not be properly captured in our simulations. The cryo-EM structure contains the stabilising mutation E1371Q, introduced to abolish ATP hydrolysis and capture CFTR in an open(-like) conformation;^12^ this residue was reverted to the wild-type (WT) glutamate to model native CFTR dynamics. Ionizable residues were modeled in their standard protonation states at physiological pH (Asp and Glu deprotonated, Lys and Arg protonated; His singly protonated), consistent with predictions by PROPKA3.1.^17^ Histidine tautomers were assigned automatically by pdb2gmx, based on the local hydrogen-bonding environment. This resulted in single protonation on NE2 in all cases. The final structure was validated with PROCHECK.^17^ Because the G551D and G1349D mutations introduce Asp residues directly adjacent to ATP at the NBD interface, we generated both protonated and deprotonated forms of these residues and evaluated their p*K*_a_ values using PROPKA3.1 on MD snapshots (see Results).

**Figure 1.**
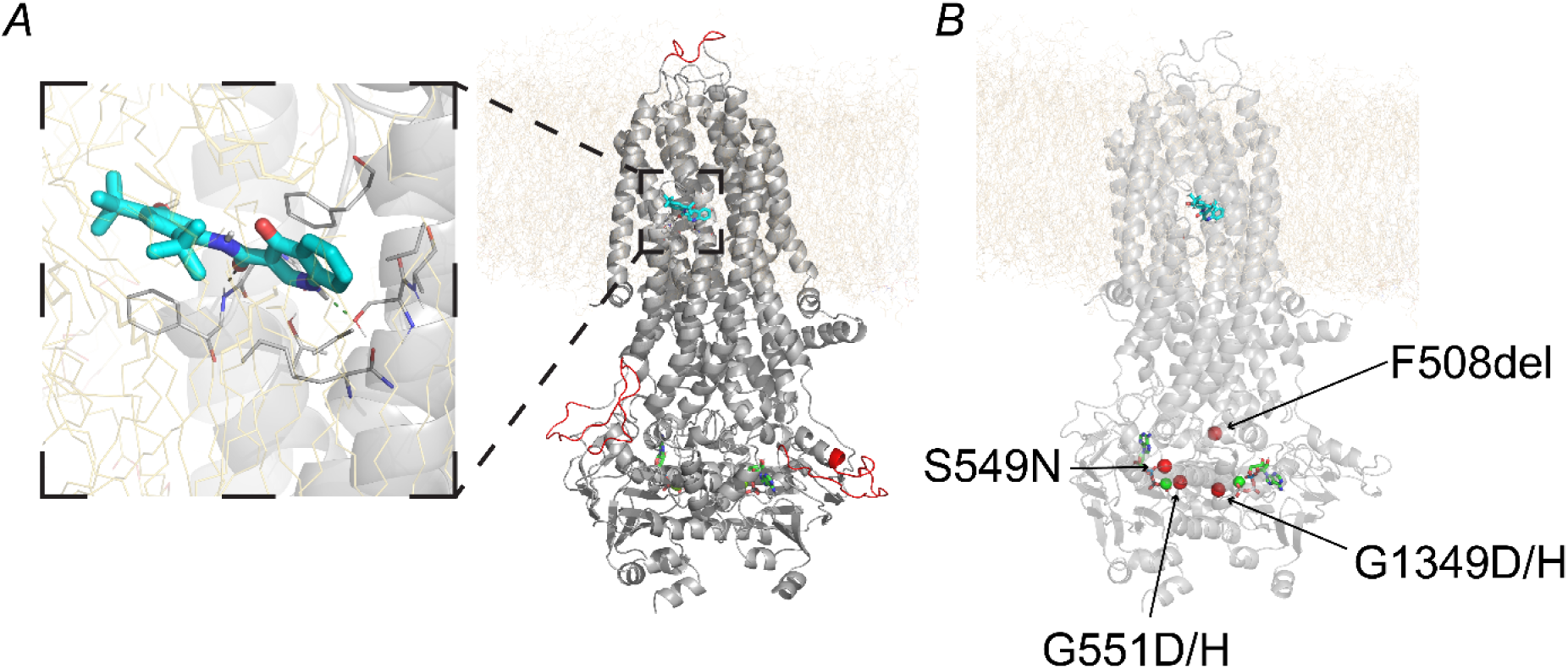
Starting structure for simulation highlighting the ivacaftor binding site and locations of studied mutations. *A*, the insert depicts a side view of the ivacaftor binding site at the protein-lipid interface. Ivacaftor is shown as sticks with cyan carbon atoms, surrounding protein residues are shown as sticks with grey carbon atoms. Hydrogen bonds between the heavy atoms of S308 O-γ and ivacaftor N16, as well as between F931 backbone N atom and ivacaftor O13, are shown as green and black dashed lines, respectively. CFTR is displayed as a grey cartoon and is embedded within a mixed lipid bilayer (POPC, POPE and cholesterol) represented by yellow sticks. ATP is shown as sticks with green carbon atoms, and Mg^2^⁺ ions as green spheres. Loops modelled from AlphaFold2 structure predictions (AF-P13569-F1-v4)^14, 15^ are coloured red. *B*, locations of CFTR mutations studied here (G551D/H, G1349D/H, F508del and S549N) are indicated as red spheres and labelled on the WT structure.

The protein, solvent, and ions were described with the AMBER99SB-ILDN force field,^19^ and the lipid bilayer with the Slipids force field.^20^ Parameters for ATP and ivacaftor were generated using generated in ACPYPE^21^ with the GAFF^22^ force field, which ensures compatibility with the force fields used for the protein and membrane. Partial charges for ATP were assigned using Gasteiger charges because higher-level schemes such as For ivacaftor, partial charges were assigned using AM1-BCC,^23^ which provides charges consistent with standard amber protein force-fields (equivalent to RESP fitting on HF/6-31G* densities) and is widely recommended for GAFF-based ligand parameterisation,^22^ especially for neutral drug-like compounds. Partial charges for ATP were assigned based on Gasteiger charges, because approaches such as AM1-BCC can strongly over-polarise phosphate groups,^24^ and can result in exaggerated ‘salt-bridge’ electrostatic interactions^25^ as found in ATP-binding sites.

We embedded the protein in a 5:5:1 POPC:POPE:cholesterol bilayer using CHARMM-GUI.^26^ The system was then solvated with TIP3P^27^ water in an 11.957 × 11.957 × 18.480 nm simulation box. Water molecules within the bilayer region (identified by *z*-coordinates relative to phosphate averages) were removed with a custom Fortran program (courtesy of Dr Richard Sessions). Following solvation and membrane drying, the system was neutralised and brought to 0.15 M NaCl. A 20 ps unrestrained *NPT* equilibration (1 fs timestep) at 200 K and 1 bar (Nosé-Hoover thermostat,^28^ 𝜏_𝑡_ = 0.5 ps; Berendsen barostat,^29^ 𝜏_𝑝_= 5.0 ps, compressibility = 4.5×10⁻^5^ bar⁻^1^) was performed to facilitate lateral compression of the bilayer, sealing the voids generated by water removal, yielding a working box of 11.38 × 11.38 × 20.78 nm.

The complete system was energy-minimized using the steepest-descent algorithm^30^ for up to 10,000 steps (or until the maximum force <200 kJ mol⁻^1^ nm⁻^1^). Direct non-bonded interactions used a 1.4 nm cutoff, and PME^31^ electrostatics (0.16 nm grid spacing, fourth-order interpolation) and a Verlet^32^ neighbour-list scheme were used.

Next, a 1 ns position-restrained *NPT* equilibration at 310 K and 1 bar (velocity-rescaling thermostat,^33^ 𝜏_𝑡_ = 0.1 ps; Berendsen barostat,^29^ 𝜏_𝑝_ = 2.0 ps, compressibility = 4.5×10⁻^5^ bar⁻^1^) was performed with a 2 fs timestep (leap-frog integrator).^34^ Bonds to hydrogen atoms were constrained with LINCS^35^ (order 4, one iteration), and non-bonded interactions used a 1.4 nm cutoff for both van der Waals and Coulomb forces. All non-hydrogen atoms of the protein were restrained with a harmonic force constant of 1000 kJ mol⁻^1^ nm⁻^2^. Velocities were regenerated at the target temperature and pressure.

To preserve the cryo-EM-identified binding position of ivacaftor while the lipid environment (which contributes ∼60% of the binding site)^9^ equilibrated, we imposed harmonic distance restraints on two key hydrogen bonds between the protein and ivacaftor (heavy atoms N of F931 with O13 of ivacaftor and O-γ of S308 with N1 of ivacaftor) using the umbrella potential implementation (𝑘 = 1000 kJ mol^−1^ nm^−2^, 𝑟_0_ = 0.28 nm) in GROMACS.^36^ These restraints were maintained throughout a 20 ns “pre-production” stage under the *NPT* ensemble at 310 K and 1 bar (Parrinello-Rahman barostat;^37^ velocity-rescale thermostat^33^). The RMSD of the ligand relative to its initial position, after alignment of the trajectory on the protein transmembrane helices, remained below 1.6 Å during the *NPT* equilibration phase. The average ligand RMSD was 1.3 ± 0.3 Å, indicating that the ligand maintained a stable binding pose throughout the simulation. After lipid equilibration, the restraints were released and unrestrained *NPT* production simulations (310 K, 1 bar) were carried out (see below), during which ivacaftor remained stably bound in the cryo-EM-defined site. To validate our membrane build, key properties were calculated using GridMAT-MD^38^ (with a 40 × 40 grid) from the WT CFTR simulations with Ivacaftor bound, based on snapshots were extracted every 20 ns from each of five independent production trajectories. The protein-corrected area per lipid (APL) was computed using the phosphate (P) atoms of POPC and POPE lipids to define headgroup positions; cholesterol was excluded because it lacks a P atom. The mean APL across 5 replicas was 52.03 ± 0.52 Å (mean ± SD, n = 5); as expected, there was no significant difference between the two leaflets. The mean (phosphate-to-phosphate) bilayer thickness was 43.2 ± 0.2 Å. The density profile (Figure S2) further reveals the expected patterns, with approximately symmetrical membrane leaflets and no water penetration in the bilayer centre.

### Molecular dynamics production runs

Production MD simulations (500 ns each) were carried out in the isothermal-isobaric *NPT* ensemble using GROMACS (v 2022.6)^39^ with periodic boundary conditions. A 2 fs integration timestep (leap-frog integrator)^34^ was used throughout. Long-range electrostatics were treated with the particle-mesh Ewald (PME) method^31^ with a 0.16 nm grid spacing and fourth-order interpolation, and a 1.2 nm cutoff was imposed for both direct-space Coulomb and van der Waals interactions (with long-range dispersion corrections). All bonds to hydrogen atoms were constrained with LINCS^35^ (order 4, one iteration). Temperature was maintained at 310 K via the velocity-rescaling thermostat^33^ (𝜏_𝑡_ = 0.5 ps), and pressure was held at 1 bar using semi-isotropic coupling with the Parrinello-Rahman barostat^37^ (𝜏_𝑝_ = 2.0 ps; compressibility = 4.5×10⁻^5^ bar⁻^1^). Trajectories were saved every 20 ps, with velocities written every 5 ns.

We simulated WT CFTR both with and without ivacaftor, as well as four mutants (Figure 1, B): the gating mutations G551D, G1349D, and S549N, and the processing mutant F508del. All point mutations were introduced in PyMOL 2.5.7^40^ on the final frame of the WT 20 ns pre-production run (with ivacaftor restraints). For the Gly-to-Asp mutations at positions 551 and 1349, both protonated and unprotonated forms were simulated (designated ’DH’ for the protonated state). The F508del construct was generated by grafting the loop from the phosphorylated, ATP- and Trikafta-bound F508del cryo-EM structure (PDB ID 8EIQ)^10^ onto the WT model. MD simulations without ivacaftor bound were prepared by deleting ivacaftor from the unrestrained production file of the respective ivacaftor-bound system, followed by 1 ns *NPT* equilibration to allow the lipid bilayer to refill the binding site. All starting structures, parameters and input files for the production runs are available via https://doi.org/10.5281/zenodo.17341486.

### Simulation analysis

Trajectories were visualised in VMD 1.9.3^41^ and PyMOL 2.5.7^40^ and quantitative analyses (RMSD, RMSF, distances, etc.) were performed using GROMACS 2022.6 tools^42^ and CPPTRAJ (part of AmberTools22).^43^ For RMSD alignment and measurement of the transmembrane helices, the following residues were selected: 81-110, 118-150, 177-218, 236-270, 292-329, 330-377, 860-880, 909-959, 966-1012, 1030-1063, 1081-1124, 1125-1162. Binding affinities of the ATP-Mg^2+^ complex to CFTR protein were estimated using molecular mechanics/generalized born surface area (MM/GBSA) calculations using the gmx_MMPBSA tool.^44^

## Results

### Exploring the structural stability of mutants in ivacaftor-free states

We calculated the root-mean-square deviation (RMSD) of C-α atoms for WT and mutant CFTR systems to assess structural stability and variations. All systems remained reasonably stable throughout the 500 ns simulations (Figure S3). To exclude the influence from the inherently flexible intracellular and extracellular loops, we also computed RMSD values after aligning the structures to the transmembrane helices (M) of the membrane-spanning domains (MSDs). These M-aligned RMSDs remained below 2 Å across all trajectories, indicating that no large-scale conformational rearrangements occurred within the transmembrane regions over this timescale. This is expected (and indicates that the model is realistic), because the timescale of our simulations is too short to capture such transitions (e.g., open-to-closed) of the CFTR channel.

### Close association of nucleotide-binding domains likely requires G551D and G1349D to be protonated

The G551D mutation lies within ATP-Binding site 2, adjacent to the negatively charged phosphates of ATP. In its deprotonated form, D551 is expected to electrostatically repel the ATP γ-phosphate, creating an unfavourable environment for binding and catalysis. To explore this, we simulated both protonation states of the D551 residue.

In initial simulations with the unprotonated (charged) form of D551, the nucleotide-binding domains (NBDs) became more separated (Figure S4), exposing D551 to solvent. In this configuration, the sidechain p*K*_a_, as calculated by PROPKA^17, 45^ on MD snapshots, was around 4, suggesting that D551 would remain charged in this conformation at physiological pH. In contrast, when D551 was protonated, the NBDs stayed closer together, creating a hydrophobic environment that shielded D551 from solvent exposure. Under these conditions, the calculated p*K*_a_ increased to ∼10, implying that D551 would be neutral in this conformation at physiological pH. This difference is reflected in our simulations (initiated from a conformation with the NBDs already close together) by a larger distance between the C-γ atom of D551 and the nearest ATP phosphate (γ-phosphate) when the residue is deprotonated (Figure 2).

**Figure 2.**
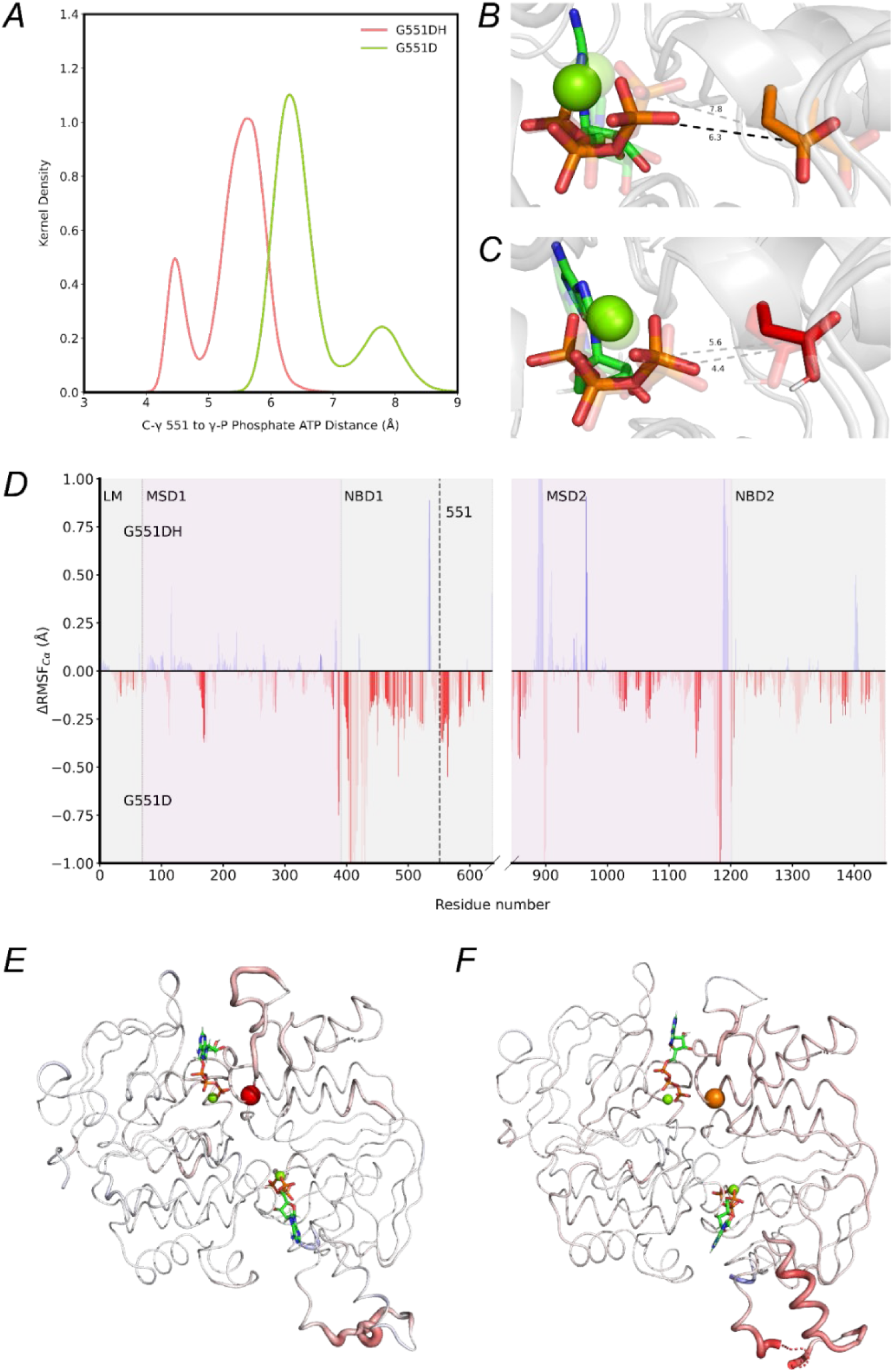
Simulation analysis of protonated (G551DH) versus deprotonated (G551D) CFTR mutant G551D. *A*, kernel-density estimates (pooled from five independent 500 ns replicates per system) of the distance (Å) between the γ-carbon (C-γ) of residue 551 and the γ-phosphate of ATP in G551D (blue) and G551DH (pink) systems. *B, C,* representative structural conformations corresponding to the two major distance peaks identified in panel A. Structures from the major peak are depicted as opaque sticks, while structures from the minor peak are shown as transparent sticks. ATP is depicted as sticks with green carbons. G551D residues are shown as sticks with orange carbons (B), and G551DH residues as sticks with red carbons (C). Dashed lines indicate measured distances, with numeric values annotated. *D*, differences in root-mean-square-fluctuation (ΔRMSF) for C-α atoms between G551DH and G551D. Blue bars indicate regions with greater flexibility in G551DH, while red bars indicate increased movement in G551D. Opacity of bars reflects statistical significance (*p* < 0.005). Residue 551 is marked by a vertical dashed line. Each CFTR domain is shaded in distinct colours and labelled above the plot. *E, F,* ΔRMSF (C-α) relative to WT mapped onto the NBDs of CFTR for G551DH (*E*) and G551D (*F*). RMSF values were mapped onto the cartoon representation as both colour and tube thickness. A blue-white-red colour scale was used in PyMOL,^40^ with red and greater thickness indicating increased flexibility compared to WT. The position of residue 551 is shown as a red sphere (E) for G551DH and an orange sphere (F) for G551D. ATP is depicted as sticks with green carbons, and Mg^2+^ is shown as small green spheres. Abbreviations: LM, lasso motif; MSD, membrane-spanning domain; NBD, nucleotide-binding domain.

Comparison of root-mean-square fluctuations (ΔRMSF) for C-α atoms revealed greater local flexibility in the G551D mutant, particularly in NBD1, compared to its protonated counterpart G551DH (Figure 2D). This enhanced mobility indicates that the charged form of D551 destabilises the NBD positioning observed in WT CFTR, leading to increased local conformational fluctuations. Similarly, when compared to WT, the charged G551D exhibits significantly higher flexibility in NBD1 (Figure 2E; Figure S5). Together, these results suggest that G551DH best represents G551D-CFTR in conformations where the NBDs adopt a conformation close to that of catalytically active CFTR,^46, 47^ with the NBDs in close proximity (Figure S4).

Simulations of the “mirror” mutation, G1349D, located at ATP-binding site 1 produced analogous results: the D1349 side chain remained closer to the ATP phosphates in its protonated state (< 7 Å) than in the deprotonated (charged) form (predominantly ∼8 Å; Figure S6). Together, these data suggest that both D551 and D1349 likely need to be protonated to facilitate closer interactions with ATP and thus a more stable conformation of the NBDs. Therefore, from this point forward, we focus exclusively on the protonated forms of the mutants: G551DH and G1349DH.

### The effects of CFTR mutations on nucleotide binding domain interactions

To assess differences in flexibility between the CFTR variants, we extended the ΔRMSF analysis (ΔRMSF = RMSF_mutant_ – RMSF_WT_) also to the remaining mutants in this study: S549N, G1349DH, and F508del (Figure S7). S549N-CFTR exhibited increased flexibility compared with WT, particularly in the NBDs, with greater fluctuations than G551DH. Although the mutation lies in the opposing NBD, G1349DH showed similar increases in flexibility in NBD1 relative to WT, and, uniquely, in MSD1 at residues corresponding to transmembrane helices M5 and M6. These local increases in NBD mobility are related to conformational changes, as quantified using principal component analysis (PCA) of the combined trajectories of all variants. the second principal component (PC2), which primarily involves NBD interdomain motions, discriminated WT from the S549N, G551DH, and G1349DH mutants (Figure 3). By contrast, F508del’s PC2 distribution and NBD ΔRMSF resembled WT, apart from elevated flexibility at residues 505-507 and 509 adjacent to the deletion site (Figure S7). Taken together, these data reveal that S549N, G551DH, and G1349DH perturb both local NBD dynamics and global inter-domain motion.

**Figure 3.**
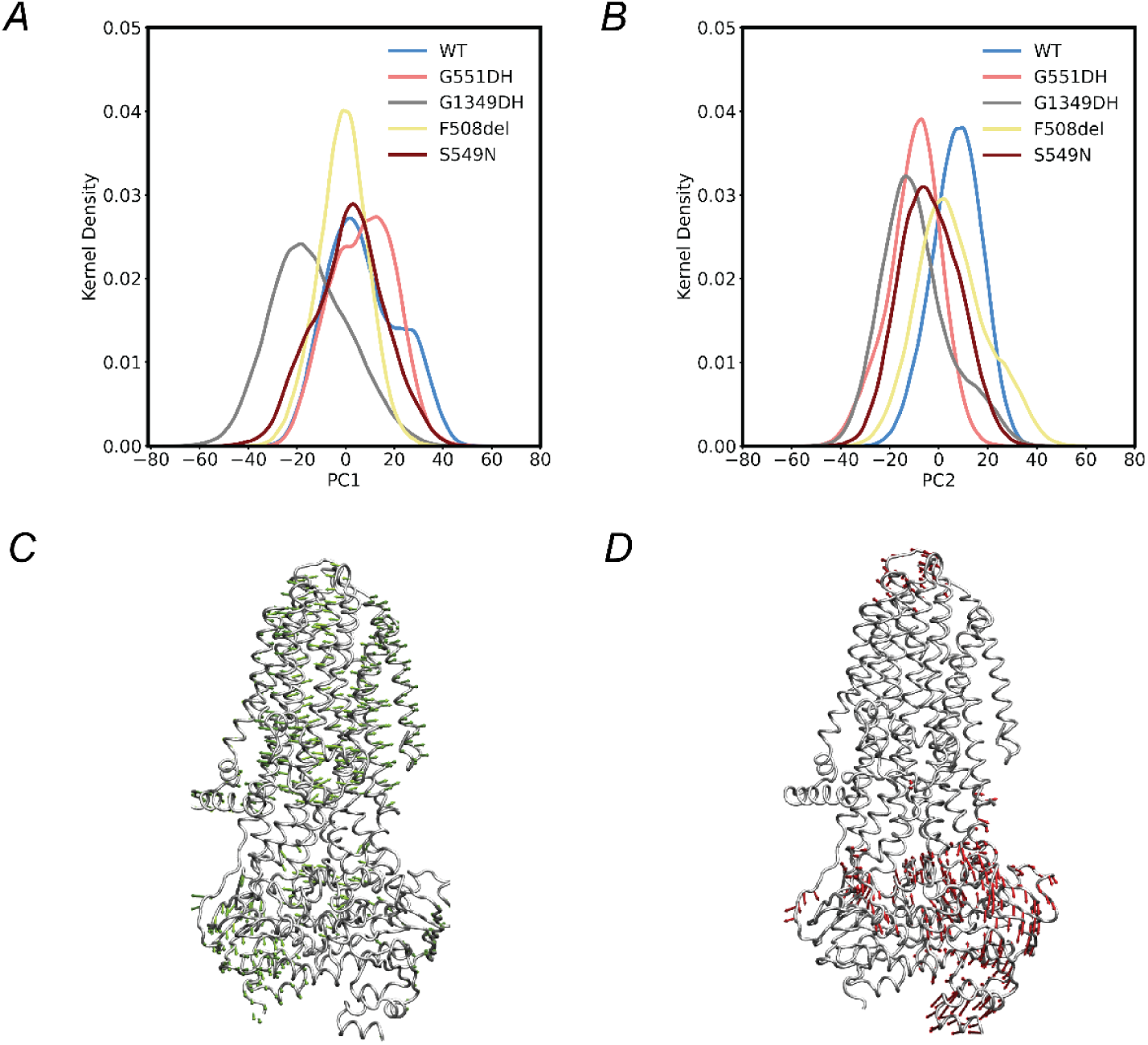
Principal Component Analysis (PCA) of WT and mutant CFTR systems. *A*, kernel-density estimate (KDE) plots (pooled from five independent 500 ns replicates per system) of the first principal component (PC1) respectively, for WT and four mutants: G551DH, S549N, G1349DH and F508del. *B*, KDE plot of the second principal component (PC2) for the same systems. *C*, *D*, conformational displacement along PC1 and PC2 mapped onto the CFTR structure. The CFTR protein is shown as a grey cartoon, and arrows indicate the principal directions of motion.

To quantify local and global differences in interactions between NBD1 and NBD2 across CFTR variants, we calculated three metrics for each system: (1) the distance between the C-α atom of residue 551 and the nearest ATP phosphate in binding site 2 (Figure 4A), (2) the distance between the C-α atom of residue 1349 and the nearest ATP phosphate in binding site 1 (Figure 4B), and (3) the centre-of-mass (COM) separation between NBD1 and NBD2, computed using only residues in α-helices and β-sheets to minimise bias from flexible loops (Figure 4C).

**Figure 4.**
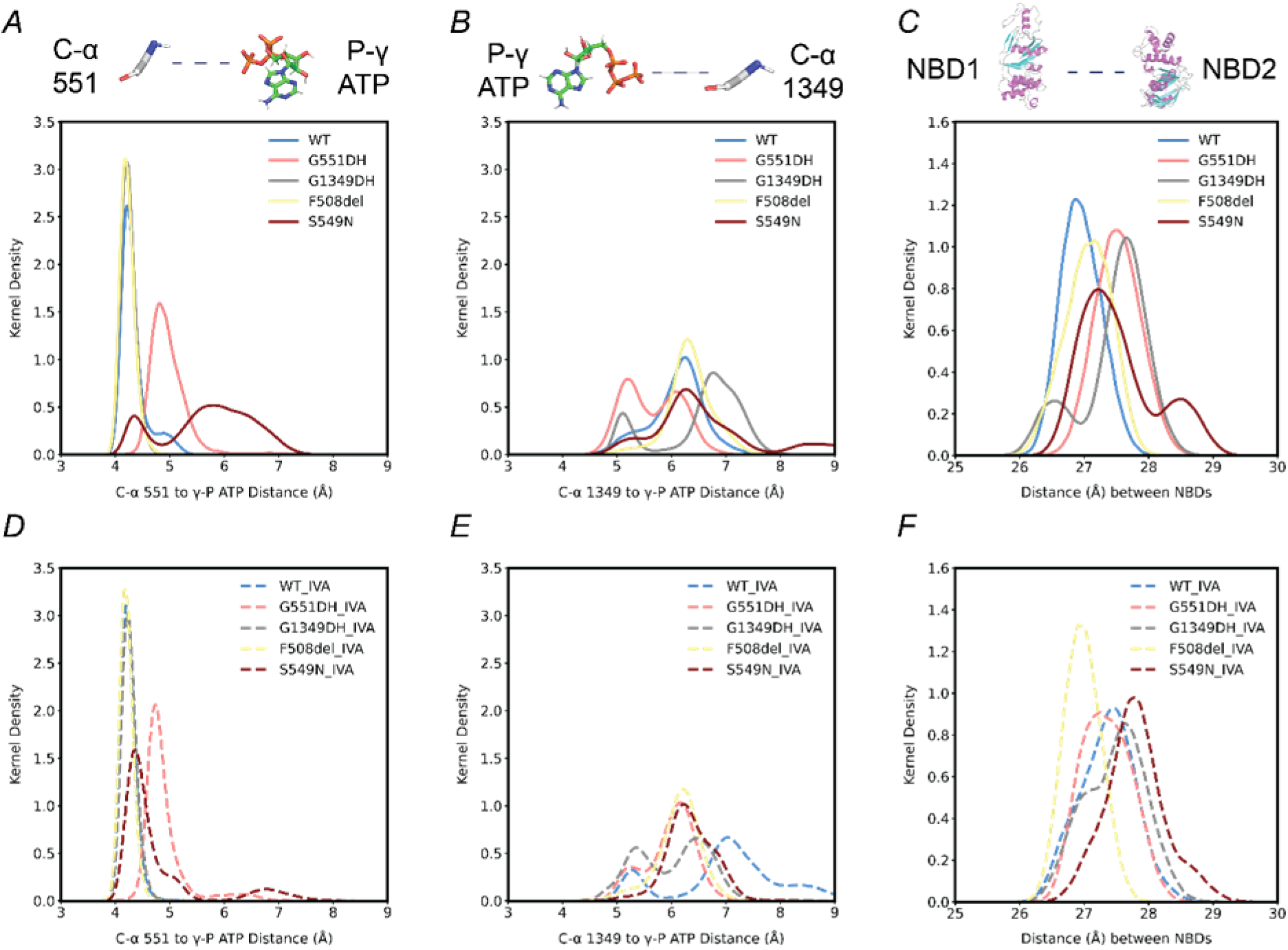
NBD inter-domain interactions in CFTR variants with and without ivacaftor, quantified by distance distributions. Kernel-density estimates (pooled from five independent 500 ns replicates per system) for unbound systems (A-C) and ivacaftor-bound systems (D-F): A, D, distance between the C-α atom of residue 551 and the γ-phosphate (γ-P) atom of ATP in the second ATP-binding site. B, E, distance between the C-α atom of residue 1349 and the γ-P atom of ATP in the first ATP-binding site. C, F, centre-of-mass (COM) separation between NBD1 and NBD2 (calculated over α-helices and β-sheets only).

For the distance between residue 551 and ATP γ-phosphate in ATP-binding site 2, WT, G1349DH, and F508del display overlapping distributions centred at ∼4 Å (Figure 4A). In contrast, G551DH shows a moderate (but significant) increase in separation (∼5 Å), and S549N displays the largest displacement and broadest range, indicative of a destabilised interaction. In ATP-binding site 1, WT and F508del again coincide, whereas G1349DH, as expected, shows the greatest separation (Figure 4B). Interestingly, a subpopulation of G551DH samples shorter distances than the main peak in WT, suggesting that this mutation affects ATP interactions in both sites. The separation between NBD1 and NBD2 is smallest for WT and only marginally higher for F508del, consistent with preserved inter-domain contacts (Figure 4C). All other mutants exhibit modestly increased separations, most pronounced in G551DH and G1349D, reflecting a weakened NBD association (S549N has a minor sub-population with the largest NBD separation).

The increase in distance between the C-α atom of residue 551 and the γ-phosphate of ATP in the G551DH and S549N mutants suggests that ATP binding at site 2 may be compromised. To investigate if this can be observed in our simulations, we performed MM/GBSA calculations on the ATP + Mg^2+^ in this site. Although such calculations are highly approximate, especially for charged ligands,^48^ we use it here as a qualitative measure of potential differences in ATP binding in different protein backgrounds, within the “open” conformational ensembles generated. Compared to WT, G551DH simulations indicate an 18 kcal/mol reduction in estimated binding energy for G551DH compared to WT, a statistically significant difference (Table S1. This reduced ATP affinity in G551DH was analysed by decomposition per residue (Figure S9) showing that S549, G550, R555 and T1246 each contribute to reducing ATP affinity in G551DH, whereas interaction with D551 itself is favourable for affinity compared to G551, likely because the longer side chain at this position allows additional contacts without strong electrostatic repulsion. However, D551 may also compromise the interactions of the nearby residues (S549, R555, T1246) with ATP, leading to a less favourable ATP coordination overall. For example, the distance between the C-ζ atom of R555 and the γ-phosphate of ATP fluctuated significantly, compared to WT (Figure S8).

For the other mutants investigated, differences in estimated ATP affinity to WT were not statistically significant (Table S1). Decomposition of residue contributions further indicated that there is essentially no effect on ATP interaction for F508del or G1349DH (Figure S10), likely because these mutations lie outside the second ATP-binding site. For the S549N mutant (which has a slightly higher estimated ATP affinity, but not significantly so; *p* = 0.43; Table S1), located within the same interaction sphere as G551DH, residues G551, Q552, K1250, and Q1291 contribute unfavourably to ATP binding (ΔΔG > 1 kcal/mol), which is partially offset by contributions from S1251 and S1255 (Figure S10). These results suggest that S549N can similarly perturb local ATP contacts as G551DH, subtly disrupting ATP positioning.

### Ivacaftor binding site is consistent across WT and mutant CFTR

We analysed the local lipid environment of the ivacaftor-binding site in WT and five mutant systems (indicated with the suffix “_IVA”). For each system, we identified the ten closest lipids of each type, namely 1-palmitoyl-2-oleoyl-sn-glycero-3-phosphocholine (POPC), 1-palmitoyl-2-oleoyl-sn-glycero-3-phosphoethanolamine (POPE), and cholesterol, and calculated the average percentage of each lipid type within 5 Å of ivacaftor (Figure 5). POPC was present most around ivacaftor, followed by POPE, then cholesterol. Visual inspection confirmed that ivacaftor contacts the lipid tails rather than the polar headgroups or ester linkages.

**Figure 5.**
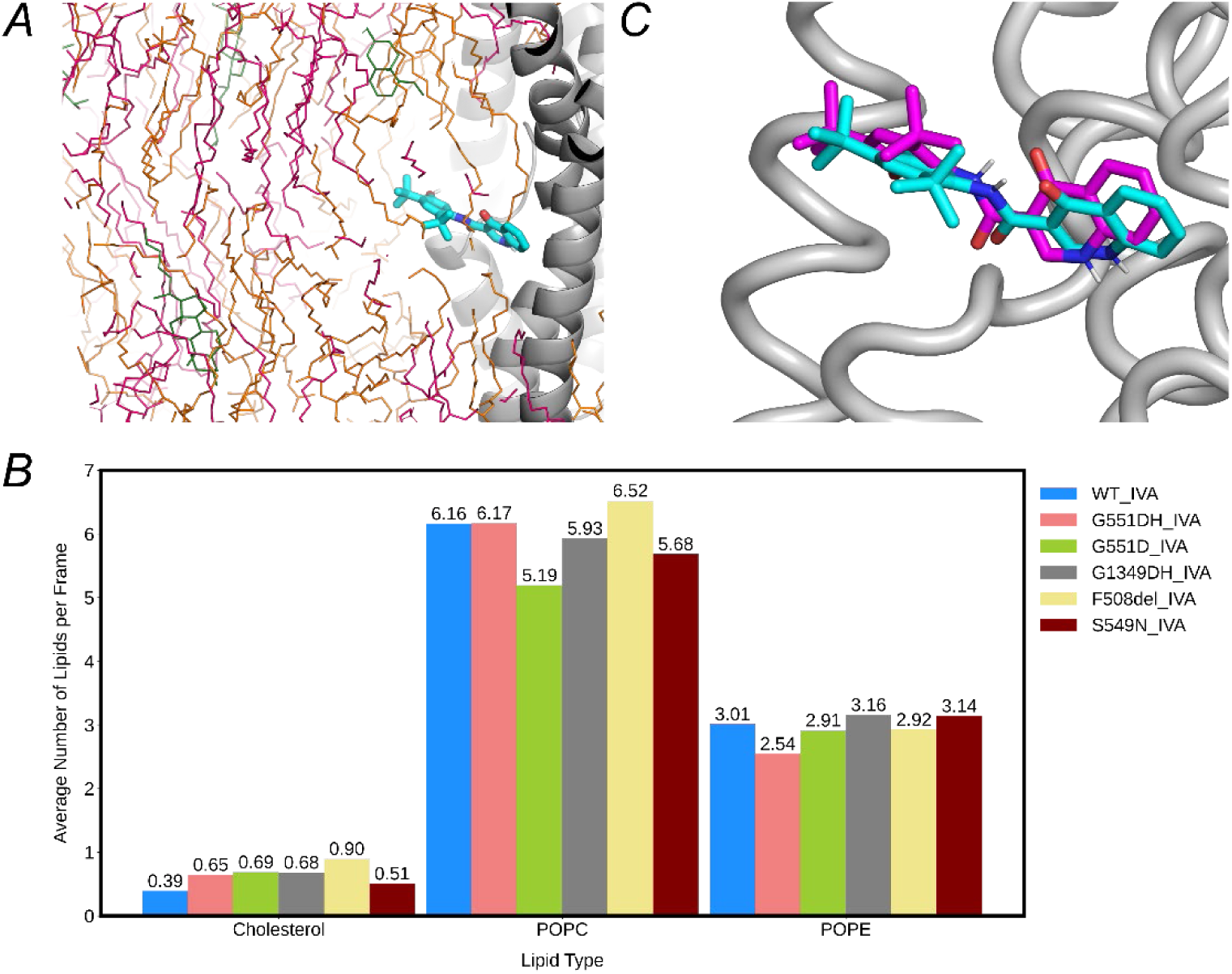
Lipid composition and binding-mode analysis of ivacaftor. *A*, Side view of ivacaftor at the protein–lipid interface (CFTR, grey cartoon; ivacaftor, cyan sticks; POPE, magenta; POPC, orange; cholesterol, green). *B*, Average number of each lipid type within 5 Å of ivacaftor per frame, pooled from five 500-ns replicates. *C*, RMSD-based *k*-means clustering (k = 3) of ivacaftor conformations, viewed from the bottom, with the mean structure (ribbon) overlaid. Trajectories were aligned on the C-α atoms of transmembrane helices (residues 81-110, 118-150, 177-218, 236-270, 292-329, 330-377, 860-880, 909-959, 966-1012, 1030-1063, 1081-1124, 1125-1162). Cluster representatives correspond to a dominant state (cyan, 56.9% of frames), a secondary state (pink, 43.1%), and a rare outlier (purple, 0.003%) not shown.

All systems showed stable ivacaftor binding (Figure S11). Hydrogen-bond analysis (data not shown) revealed that two key interactions (see Figure 1A) are conserved across all variants: between S308 O-γ and ivacaftor N16 (present in ∼80% of frames, based on a maximum donor-acceptor distance of 3.0 Å and a minimum donor-H-acceptor angle of 135°), and between F931 N and ivacaftor O13 (present in ∼60% of frames). Ivacaftor also adopted comparable conformations in all simulations (Figure 5C). RMSD-based k-means clustering after alignment on the C-α atoms of transmembrane helices, revealed two slightly different conformational states, representing 56.9% and 43.1% of the sampled frames. Frame distributions among these clusters were virtually identical for WT and all mutants (Table S2). Overall, therefore, ivacaftor binding in our simulations is essentially the same between WT and mutant CFTR systems.

### Ivacaftor’s influence on the NBDs and ATP binding is mutation dependent

To determine the impact of ivacaftor (IVA) on NBD inter-domain interactions in CFTR, we applied the same three metrics as described above (Figure 4). In WT, ivacaftor does not alter the 551-to-ATP distance, but it causes an increase in both the 1349-to-ATP distance and the NBD1-NBD2 distance compared to the drug-free state. In G551DH, ivacaftor also does not substantially alter the 551-to-ATP distance (distribution similar in drug-free and drug-bound states). However, the NBD distance shifts closer to that of drug-free WT, possibly indicating a partial rescue of NBD association. In G1349DH, ivacaftor appears to reduce the 1349-to-ATP distance towards the drug-free WT value, while the NBD distance remains similar to before. This suggests that ivacaftor might primarily improve the local interaction at ATP-binding site 1 without significantly affecting overall NBD separation. In S549N, ivacaftor partially restores the 551-to-ATP distance towards the drug-free WT conformation (with limited change to the 1349-to-ATP distance). In contrast, the NBD distance increases relative to both unbound S549N and unbound WT, indicating a more complex alteration in the NBD interface that may reflect incomplete or differential rescue of NBD association in this mutant. In F508del, the metrics are similar to those of drug-free F508del (and drug-free WT), indicating that ivacaftor does not significantly affect NBD association or ATP-binding site geometry in this variant, which was already similar to WT.

Notably, the predicted ATP binding affinity at the second ATP-binding site in G551DH with ivacaftor bound was no longer significantly different compared to WT-CFTR, compared to without ivacaftor (Table S1). Although this suggests partial rescue of ATP binding by ivacaftor, the difference in binding affinity between ivacaftor-bound and ivacaftor-free G551DH did not reach statistical significance (*p* > 0.1). Thus, while ivacaftor might improve ATP binding compared to untreated G551DH, it does not fully restore it to WT levels. Residue-level energy decomposition (ΔΔG) further elucidates these findings. Comparing G551DH_IVA with drug-free WT revealed no significant change in R555’s contribution to ATP binding (Figure S14), whereas drug-free G551DH did show a change. However, S549, F550 and T1246 remained destabilising, mirroring the pattern seen in drug-free G551DH. In a direct comparison of G551DH_IVA to G551DH (that is, ivacaftor-bound mutant versus ivacaftor-free mutant), only residue S549 exhibited a statistically significant difference. Taken together, these results indicate that ivacaftor may confer some improvement in ATP binding for G551DH, but underlying residue-level interactions, particularly around S549, F550, and T1246, still deviate from the WT, consistent with only partial restoration.

### The relationship between ivacaftor binding and CFTR pore interactions

Two chloride pore salt bridges, R347-D924 and R352-D993, have been identified as crucial for maintaining the open-pore architecture of CFTR,^49, 50^ which are expected to be stable in WT CFTR (leading to long, stable channel openings^46^). In our simulations, we monitored these putative salt bridges through the distance between the functional-group carbon atoms (C-ζ to C-γ), using a threshold of 6 Å to define a “formed” salt bridge. Our results show that the R347-D924 salt bridge maintains a distance of 4-5 Å throughout the simulations (>95% of the frames, consistent with the cryo-EM structure used as starting point of the MD simulations) for both WT and mutant systems (Figure S15). In contrast, the R352-D993 interaction is only formed intermittently, present in about 30% of frames in WT and even less in some mutants (7% in F508del and G1349DH; Figure 6). Time-resolved analyses show repeated formation and breakage throughout the trajectories (data not shown). Notably, ivacaftor appears to significantly increase the R352-D993 salt bridge occupation in G551DH and F508del.

**Figure 6.**
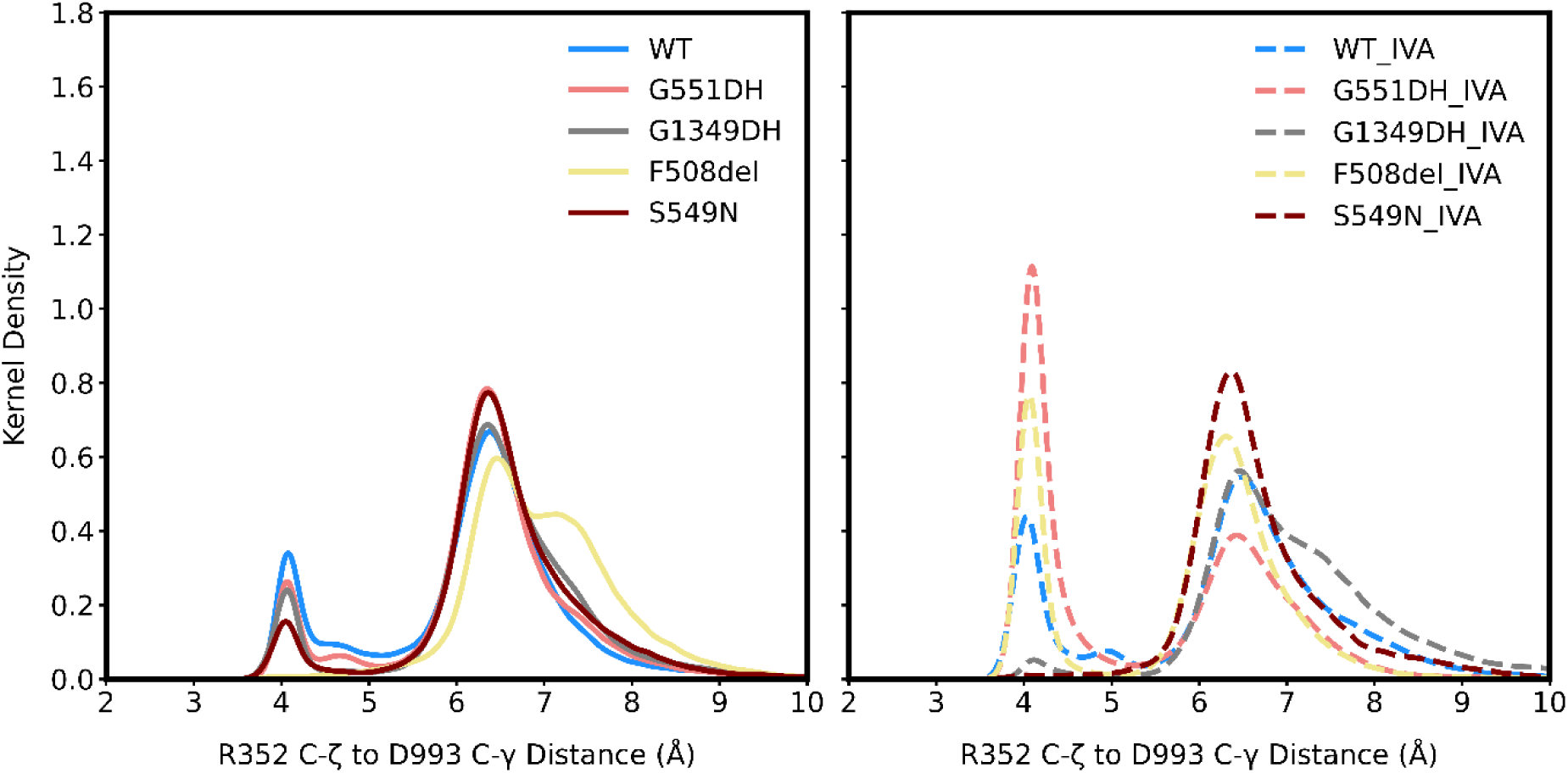
Pore-domain salt bridge R352–D993 formation in CFTR variants, quantified by distance distributions. Kernel-density estimates (pooled from five independent 500 ns replicates per system) of the distance between the C-ζ atom of R352 and the C-γ atom of D993 for simulations without ivacaftor (left) and with ivacaftor (right).

### Water passes through the CFTR pore in simulation but not chloride ions

The CFTR pore has an hourglass shape, with a shallow outer vestibule and a deep cytosolic vestibule separated by a narrow constriction, referred to as the selectivity filter.^9, 10, 51^ Recent work from Levring and Chen^52^ proposes that a hydrated chloride (Cl^−^) ion enters via a lateral portal between transmembrane segments M4 and M6, where it is stabilised by a positive electrostatic surface. As the Cl^−^ ascends, it dehydrates to pass through the selectivity filter, where it is stabilised by interactions with G103, R334, F337, T338, and Y914. Upon moving through a narrow lateral exit between M1 and M6, the Cl^−^ then rehydrates as it enters the epithelial lumen. In complementary microsecond-scale MD simulations conducted under a strong hyperpolarising electric field, spontaneous Cl⁻ permeation occurred in 2 of 10 runs, accompanied by subtle M1 and M6 rearrangements.^53^ However, equivalent simulations without the electric field showed Cl⁻ binding in the channel pore, but no permeation.

We investigated chloride-CFTR interactions in our simulations of WT and mutant CFTR. For this purpose, the membrane region was defined by the phosphate-group *z*-coordinates and any ion entering this portion was followed. To focus on sustained, physiologically relevant events, only ions present in more than 10% of the frames were included in the analysis. In all our simulations, a single Cl⁻ occupies the pore for the majority of the time, although the specific ion frequently swaps (Figure 7, Figure S16; the latter shows chloride occupancy of the pore for each simulation replica). Consistent with previous work on WT CFTR^53^, we did not observe a Cl⁻ crossing the upper membrane boundary in any of our WT or mutant CFTR simulations, indicating that the pore does not permit spontaneous ion movement into the extracellular space at this timescale (without applying an electric field). Levring and Chen suggest that positively charged residues attract Cl⁻ from the intracellular (bottom) side toward the extracellular (top) side of the pore,^52^ a phenomenon that we also observed across these CFTR systems. In our simulations, Cl⁻ ions are observed to interact with R334, a residue located within the selectivity filter, indicating an upward attractive force (Figure 7).

**Figure 7.**
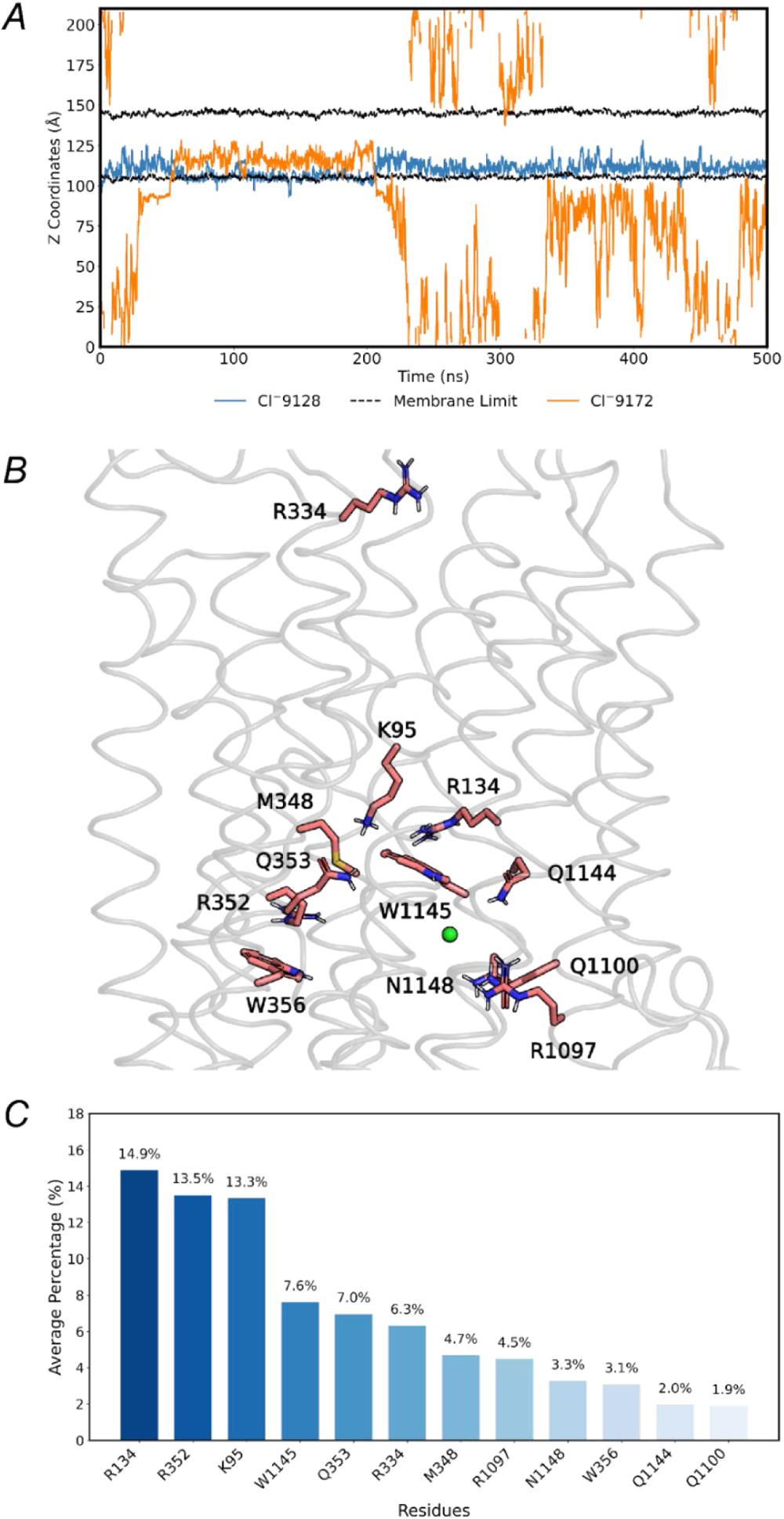
Chloride-ion occupancy and residue interactions in the CFTR pore. *A*, time-resolved *z*-coordinate positions of Cl⁻ ions entering the membrane region (defined by the phosphate-group *z*-coordinates) during one 500-ns WT CFTR simulation (one of five independent replicates). Two individual ions are highlighted in blue and orange and the upper and lower membrane boundaries are shown as black dashed lines. *B*, CFTR pore residues that contact Cl⁻ in > 1% of frames are shown as pink sticks on the grey cartoon backbone; a representative pore Cl⁻ is shown as a green sphere. Data are pooled across all systems, with and without ivacaftor, over five independent 500-ns replicates each. *C,* mean percentage of frames in which each common residue interacts with Cl⁻ (≤3.5 Å), pooled across all systems ± ivacaftor (five 500-ns replicates each).

Across all systems, we quantified interactions between Cl⁻ ions and CFTR residues by identifying any pore residue with at least one atom within 3.5 Å of Cl⁻ for more than 1% of the simulation frames. Notably, aside from the aforementioned R334, these interactions consistently involved residues Q353, W1145, Q1100, R352, R134, K95, N1148, R1097, Q1144, M348, W356, all located around the inner vestibule of the channel pore (Figure 7).

We applied the same tracking approach for water molecules as used for Cl^−^, observing frequent, bidirectional water movement through the CFTR pore in our simulations (Figure S17). The number of such passages differs per simulation, but reaches several hundred in each. There is no clear evidence of ivacaftor (or mutations) altering the rate or extent of water passage (Figure S18). Thus, the mutations and ivacaftor appear to have minimal impact on water permeability, or at least that any such effect is not readily discernible from our current simulations.

## Discussion

In this work, we present extensive all-atom molecular dynamics (MD) simulations of the membrane-embedded cystic fibrosis transmembrane conductance regulator (CFTR) in both wild-type (WT) and disease-causing mutant forms, and either with ivacaftor bound or not. Using 32.5 µs of MD simulations (13 systems, each with five independent 500 ns repeats), we investigated how CF-associated mutations, including gating mutations G551D, G1349D and S549N, and the folding/gating F508del mutation, alter CFTR structure and dynamics. While individual 500 ns simulation runs may not capture large-scale conformational rearrangements away from the starting model, they are well suited to reveal local structural and flexibility changes relevant to CFTR function.

In WT CFTR, the binding of one ATP molecule (coordinated by a Mg^2+^ ion) at each of the two ATP-binding sites drives close association of the two nucleotide-binding domains (NBD), a process commonly referred to as NBD dimerisation.^54, 55^ In the dimerised state, each ATP molecule simultaneously engages the Walker A (phosphate-binding) motif of one NBD and, from the opposing NBD, the ABC signature sequence (LSGGQ in NBD1 or its degenerate LSHGH in NBD2).^55^ Gating mutations G551D and G1349D both substitute Gly with Asp within these signature motifs (resulting in LSGDQ in NBD1, or LSHGH in NBD2, respectively). G551D is delivered to the cell surface at levels comparable to WT^56^ but fails to activate the chloride channel.^57^ In WT CFTR, the backbone amide nitrogen of G551 lies within hydrogen-bond distance of the ATP γ-phosphate in the dimerised state.^58^ Replacing Gly with Asp could introduce a negative charge, that likely repels the γ-phosphate. Under saturating ATP (3 mM), single-molecule FRET between NBDs indicates an intermediate FRET efficiency for G551D (∼0.37), distinct from both fully dimerised WT (∼0.49) and the NBD-separated CFTR (∼0.25), suggesting a disrupted but not abolished NBD approach.^47^ Our MD simulations start from the fully dimerised CFTR conformation. When Asp is treated as unprotonated (negatively charged), the distance between residue 551 and the nearby γ-phosphate is increased compared to WT. However, with Asp singly protonated (consistent with the expected p*K*_a_ at the dimerised conformation), the distance only increases by ∼1 Å compared to WT CFTR. The potentiator ivacaftor appears to have minimal impact on this interaction, inducing at most a very slight reduction in the D551 to γ-phosphate distance. Our simulations suggest that with Asp unprotonated, NBD dimerization is not stable, and indicate the start of NBD separation. Therefore, we suggest that two states of G551D CFTR exist under physiological conditions: one (partially) NBD-separated, inactive, conformation with D551 unprotonated, and one NBD-dimerised state, with D551 protonated. Once the NBDs approach sufficiently (with ATP bound), the local microenvironment would shift the p*K*_a_ such that D551 becomes protonated, and full NBD dimerization can occur. Nevertheless, even when D551 is protonated, its interaction with ATP does not fully recapitulate the local NBD-dimerized conformation observed in WT CFTR, implying additional effects beyond simple charge repulsion. Indeed, binding affinity predictions confirm that ATP in NDB2 is bound with lower affinity with protonated D551 (G551DH) compared to WT. Recent cryo-EM data on G551D in complex with ivacaftor support the idea that ivacaftor enhances CFTR function by increasing the coupling efficiency between NBD dimerisation and pore opening, thereby prolonging the open state and enhancing chloride flux.^59^ Although ivacaftor does not primarily promote NBD dimerisation, we propose that it may exert a secondary effect by helping to stabilize the NBD interface, therefore reducing the solvent exposure of D551, shifting its equilibrium towards the protonated state. Once protonated, productive interactions with ATP in binding site 2 can be maintained, as indicated by our simulations.

G1349D mirrors G551D in NBD2’s (degenerate) signature motif.^57^ Unlike G551D CFTR, which is largely ATP-insensitive, G1349D retains some ATP dependence but lowers the channel’s open probability (P_o_) approximately tenfold versus WT.^57^ Although ivacaftor can potentiate G1349D CFTR (increase in P_o_), genistein, another known CFTR potentiator that stimulates both WT and F508del-CFTR, does not do so.^60^ Our simulations suggest that, as for G551D, protonation of D1349 is likely required to mitigate electrostatic repulsion with the ATP phosphate groups in the NBD-dimerised conformation. When protonated (G1349DH), the ATP binding to NBD2 (near G551) is essentially equal to WT CFTR, as is the flexibility of NBD2, where the mutation is located. However, increased flexibility in transmembrane segments M5 and M6 observed in our simulations supports the existence of an allosteric pathway between the NBDs and the chloride pore that may underlie G1349DH’s defective gating. Notably, ivacaftor reduces the distance between the protonated residue D1349 and ATP, restoring it towards WT values. This may indicate that interactions between ATP in NBD1 and the region with the mutation are strengthened, which in turn may contribute to the G1349D CFTR potentiation (and thus better CFTR function).

The F508del mutation in CFTR destabilises the protein folding, triggering endoplasmic reticulum quality-control recognition and leading to degradation before the channel can traffic to the cell surface.^61^ Even when pharmacologically rescued to the plasma membrane,^62^ F508del CFTR displays gating defects that require potentiator treatment. In this study, we observed that conformational dynamics and ATP binding of F508del CFTR were similar to WT, but noted a reduction in MSD1 flexibility that may hinder the structural transitions required for ion permeation. Ivacaftor binding increased flexibility in both MSD1 and NBD1, which could underlie its potentiating effect by facilitating gating transitions, but can also explain why chronic treatment of F508del-carrying patients with ivacaftor can destabilize F508del CFTR.^63^

S549 lies within the conserved LSGGQ signature motif of NBD1, where its side chain hydroxyl normally hydrogen-bonds with the ATP γ-phosphate.^12, 16^ Our simulations indicated that the mutation disrupts this interaction, shifting the γ-phosphate further from position 549 and compromising the alignment needed for hydrolysis. Functionally, S549N is classified predominantly as a gating mutation,^16^ with some mild processing defects.^64^ ATP binding affinity and functional efficacy (as measured by current passage) of S549N are diminished compared to WT.^64^ When treated with ivacaftor and lumacaftor (a corrector that mitigates processing defects), S549N currents recover to near-WT levels. In our simulations, the γ-phosphate of ATP is indeed further from residue 551 in S549N compared to both WT and G551DH. Interestingly, with ivacaftor present, this distance is partially “rescued” to WT levels. Notably, despite these local improvements in ATP binding, ivacaftor increases the overall NBD distance somewhat, suggesting a nuanced allosteric effect. This supports the notion that ivacaftor does not directly promote increased gating in CFTR by causing closer NBD association, but through other dynamic effects (e.g., affecting the pore). Although the S549N mutation affects the ATP local environment, our binding affinity calculations indicate that it likely does not prevent ATP binding. However, the disruptions in catalytically required alignment and increased flexibility in NBD1 that we observe indicate that conformations required for efficient ATP hydrolysis are not easily maintained.

Beyond the specific variants simulated here, our results may help rationalise the behaviour of other mutations located in or near the ATP-binding signature motifs. Clinically important mutants such as G551S and S549R, which affect the same LSGGQ signature loop in NBD1 as G551D and S549N, are similarly classified as class III gating variants^65^, causing severe defects in ATP-dependent channel gating while retaining substantial cell-surface expression, and they are responsive to potentiators such as ivacaftor *in vitro.*^56^ In NBD2, G1244E lies close to the degenerate signature motif, likely disrupts ATP interactions (due to charge repulsion between the carboxylate and ATP phosphates), and similarly produces a marked gating defect that is only partially rescued by ivacaftor. Given that all signature-motif mutants studied here (G551D, S549N and G1349D) primarily perturb productive ATP coordination and NBD coupling, rather than abolishing ATP binding outright, our simulations support a common mechanism, where other class III variants at the signature motifs, such as G551S, S549R and G1244E, similarly weaken NBD-NBD interactions and ATP-driven gating, while remaining amenable to rescue by potentiators such as ivacaftor.

Although our simulations showed that (at least) one chloride ion remains within the pore inner vestibule throughout, the ion did not pass through the selectivity filter to reach the extracellular side. Conversely, bidirectional water passage through the channel was observed in all systems. This observation suggests that additional conformational changes are required for chloride conduction, and these transitions are not captured within our simulation timescales. This aligns with previous studies suggesting that CFTR undergoes complex structural rearrangements to transition between permissive and non-permissive states for chloride conduction.^52^ The presence of ivacaftor in our simulations affected neither chloride nor water passage. We identified several key residues that consistently interact with chloride ions, including several positively charged residues that were previously indicated to be important for anion conduction, such as R352,^66^ R134,^67^ K95.^66,67^

By conducting extensive MD simulations on our (Cryo-EM-based) CFTR model in the open-channel conformation, we gained new insights into how several CF-associated mutations can subtly disrupt channel dynamics and (thereby) function, and how the modulator ivacaftor influences (and partially rescues) these processes. Importantly, our extensively tested model provides a starting point for future investigations into CFTR dynamics, which may guide the development of next-generation therapeutic strategies. To facilitate such investigations, we make our starting models and simulation protocols openly available. In this study, we observed that mutations in the ATP-binding sites such as G551D, S549N and G1349D disrupt local ATP-binding and NBD interactions, with protonation of aspartate in G551D and G1349D being a critical factor for maintaining the functionally required NBD dimerisation. We further found that ivacaftor’s primary mechanism of action likely involves modulating the local environment of the transmembrane helices rather than directly tightening NBD association. Together, these insights provide a more nuanced understanding of CFTR dynamics and the molecular mechanisms underlying its mutation- and potentiator-induced functional alterations.

## Acknowledgements

We thank Dr. Richard Sessions for his insightful discussions and invaluable coding assistance throughout this work. We also gratefully acknowledge the high-performance computing resources provided by the Advanced Computing Research Centre; HPC time on the ARCHER2 UK National Supercomputing Service (https://www.archer2.ac.uk) allocated via the UK High-End Computing Consortium for Biomolecular Simulation (HECBioSim, http://hecbiosim.ac.uk), supported by EPSRC (grant no. EP/X035603/1); and the Isambard 3 Tier-2 HPC Facility, hosted by the University of Bristol and operated by the GW4 Alliance (https://gw4.ac.uk), funded by UK Research and Innovation in conjunction with EPSRC (EP/X039137/1). DFV’s PhD studentship was funded by the Cystic Fibrosis Trust.

## Competing interests

The authors declare no competing interests.

## Author contributions

Diana-F. Veselu – conceived experiments, conducted all simulations and simulation analyses, data interpretation, initial draft writing, manuscript revision.

Deborah K. Shoemark – conceived experiments, assisted with conducting simulations and analyses, data interpretation, manuscript revision, supervision.

Marc W. van der Kamp – conceived experiments, data interpretation, manuscript writing, manuscript revision, supervision.

## Data availability

Starting structures for simulations, GROMACS topology and parameter files, and GROMACS MD input files are available at DOI: 10.5281/zenodo.17341486. All further relevant data are within the manuscript and the Supplementary Information.

## Notes

https://github.com/dianaveselu/CFTR/blob/main/bidirectional_water_passage.avi?raw=true.

### Competing Interest Statement

The authors have declared no competing interest.

### Summary of Updates

This revised version incorporates minor changes made during peer review, including additional analyses of membrane properties derived from the molecular dynamics trajectories (e.g., area per lipid, bilayer thickness and density profiles). These updates clarify the membrane environment without altering the overall results or conclusions. The revision also updates the preprint record following publication in a peer-reviewed journal.

