## Supplemental Information for "Influence of Disease-Causing Mutations and Ivacaftor on the Dynamics of the Cystic Fibrosis Transmembrane Conductance Regulator (CFTR) Protein"

### for

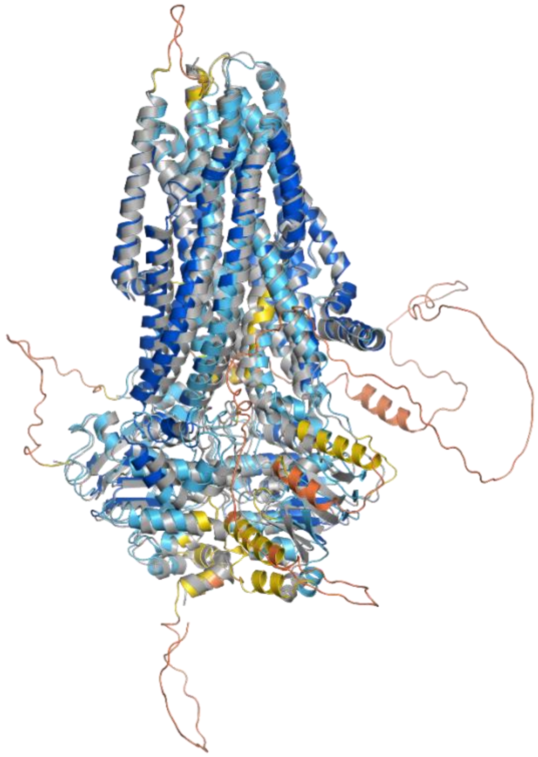

**Figure S1. Alignment of the AlphaFold2 model of phosphorylated, ATP-bound CFTR with the cryo-EM structure of the phosphorylated, ATP-bound, ivacaftor-bound CFTR (PDB ID 6O2P).**

The AlphaFold2 AF-P13569-F1-v4 model is coloured by the standard pLDDT confidence palette (red: pLDDT < 50; orange: 50-70; yellow to cyan: 70-90; blue: ≥90), while the cryo-EM structure (PDB ID 6O2P)[1] is shown as a grey cartoon.

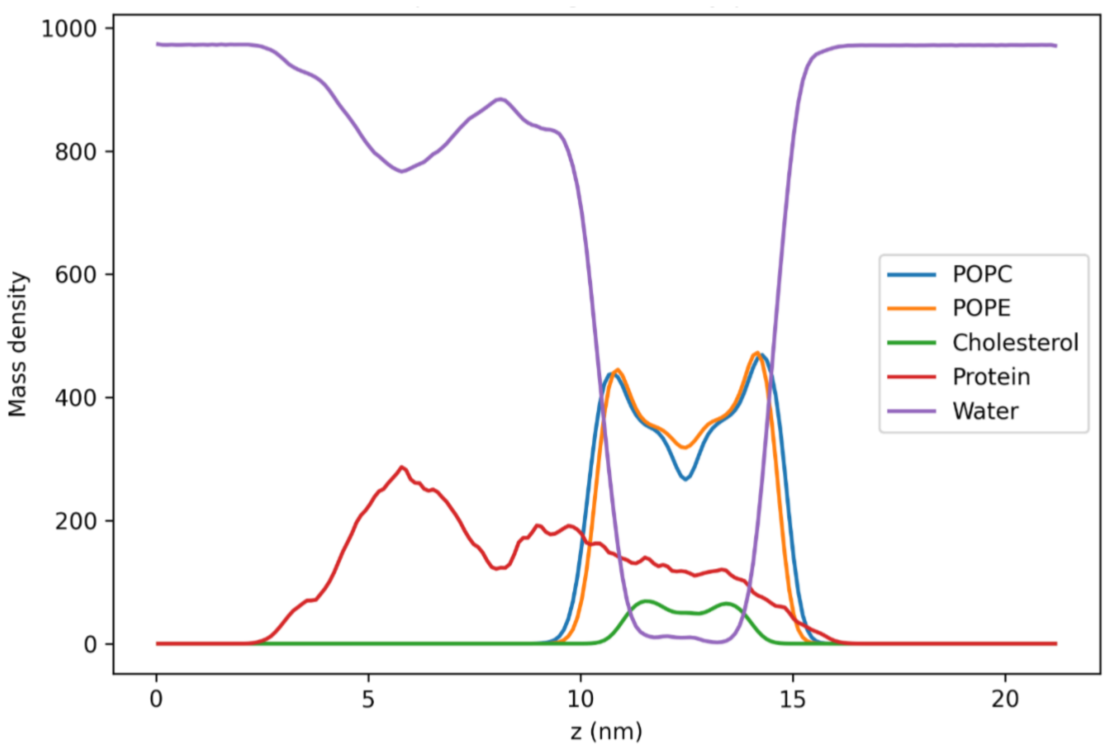


**Figure S2. Density profile for WT CFTR with ivacaftor.**

Mass density profiles along the bilayer normal (*z*-axis) for POPC (blue), POPE (orange), cholesterol (green), protein (red), and water (purple), calculated from WT CFTR simulations with ivacaftor bound. Profiles were obtained using GridMAT-MD from snapshots extracted every 20 ns across five independent 500 ns production replicas and averaged over all trajectories. The profile reveals the expected (approximately) symmetrical bilayer for the simulated 5:5:1 POPC:POPE:cholesterol mixture, with essentially no water present amongst the lipid tails.

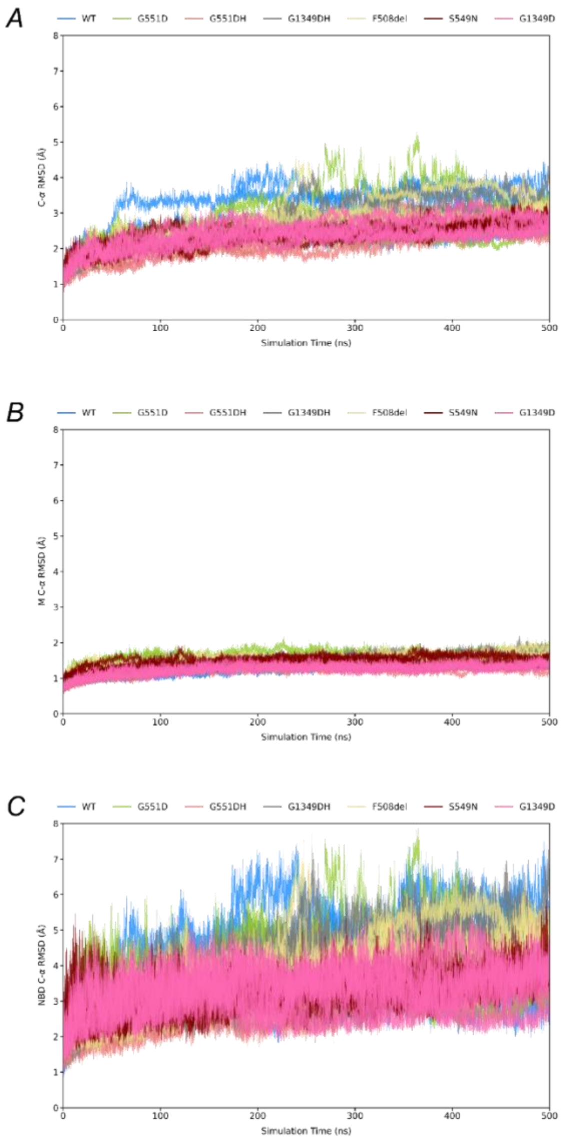


**Figure S3. Time evolution of C-α RMSD for WT and mutant CFTR systems.**

The C-α RMSD over 500 ns is shown for WT (blue) and six mutant systems: G551D (green), G551DH (red), G1349DH (grey), F508del (khaki), S549N (maroon), and G1349D (magenta). *A*, C-α RMSD calculated over all protein residues; *B*, C-α RMSD calculated over transmembrane-helix residues M1-M12: residues 81-110,118-150,177-218, 236-270, 292-392, 330-377, 860-880, 909-959, 966-1012, 1030-1063, 1081-1124, 1125-1162, as defined in Hwang *et al.*[2] with the exception of M8 (909-959) and M9 (966-1012); *C*, C-α of the two nucleotide-binding domains (NBD1: residues 391-637; NBD2: residues 1175-1452), each aligned to the transmembrane helices. Individual RMSD traces for all replicate simulations of each system are shown.

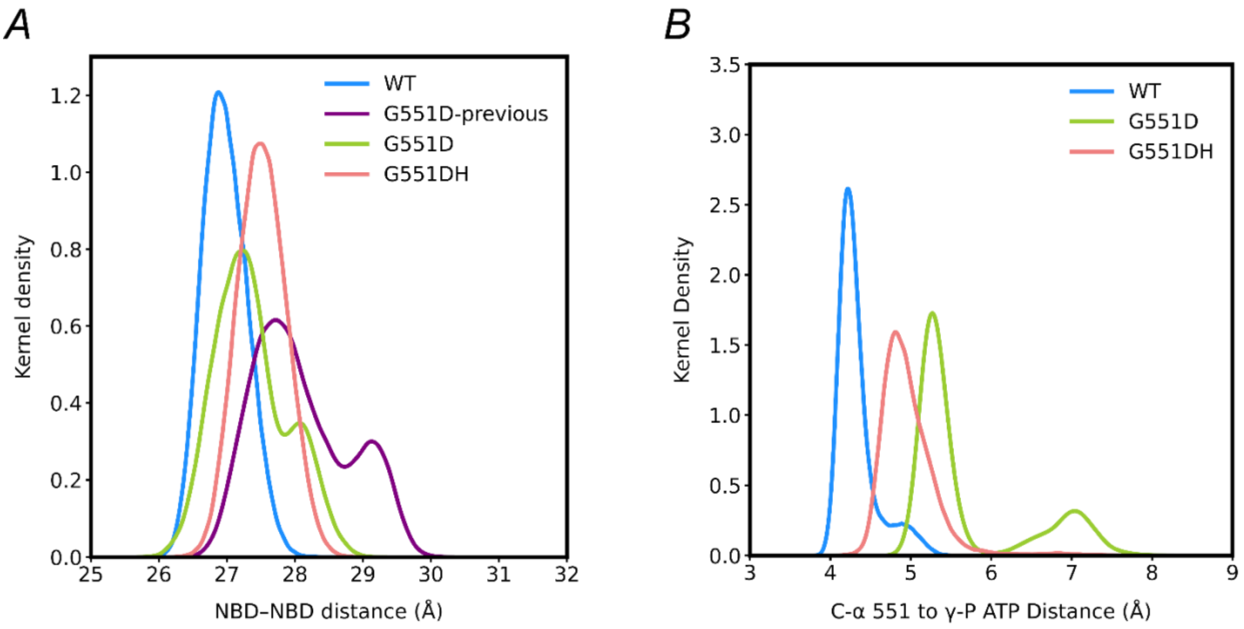


**Figure S2. Effects of D551 protonation on NBD association and ATP-site geometry in G551D CFTR.**

*A,* Kernel-density estimates (pooled from five independent 500 ns replicates per system) of the centre-of-mass separation between the α-helices and β-sheets of NBD1 and NBD2, comparing WT CFTR (blue), previous G551D simulations (purple) and new simulations in which D551 is deprotonated (green) or protonated (G551DH; pink). *B*. Kernel-density estimates (pooled from five independent 500 ns replicates per system) of the distance between the C-α atom of residue 551 and the γ-phosphate (γ-P) atom of ATP in the second ATP-binding site, comparing WT CFTR (blue), G551D (green), and protonated G551D (G551DH; pink).

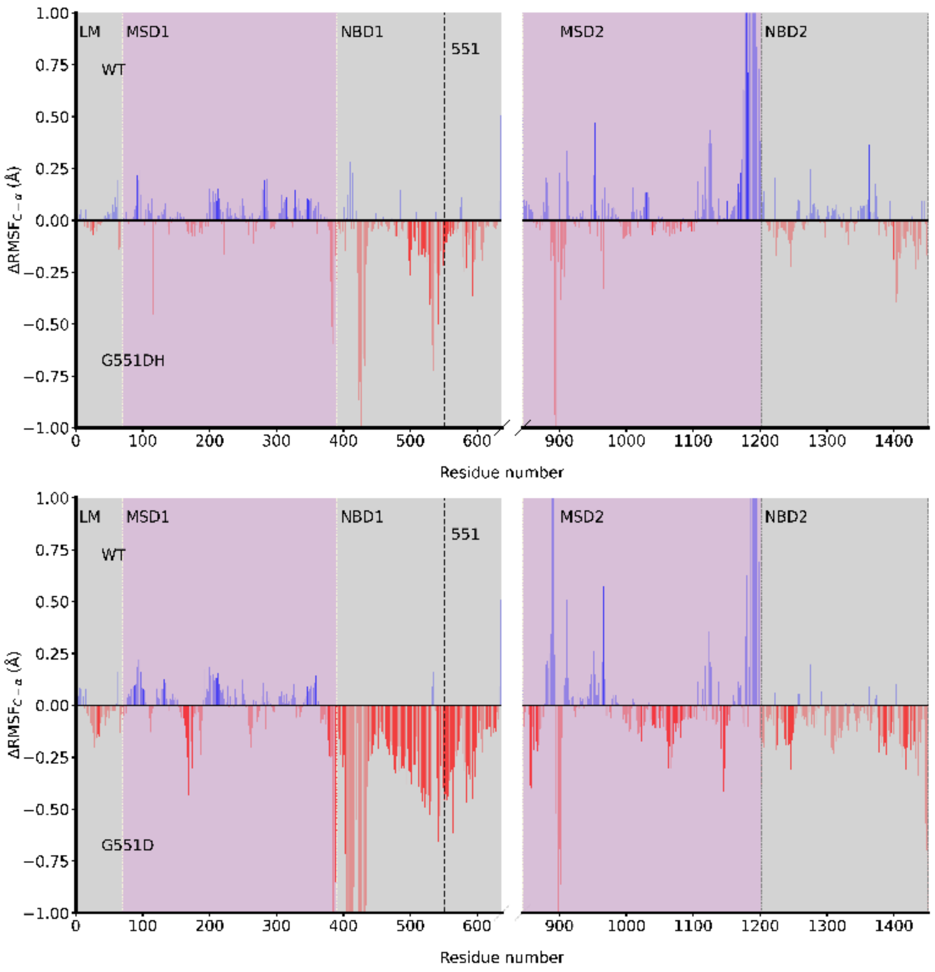


**Figure S3. ΔRMSF analysis between WT and protonated and deprotonated G551D-CFTR.**

Differences in root-mean-square fluctuation (ΔRMSF = $RMSF_{mutant} - RMSF_{WT}$) for C-α atoms comparing WT CFTR with protonated (G551DH; top panel) and deprotonated (G551D; bottom panel) states of the G551D mutant. Blue bars represent regions with greater flexibility in WT, while red bars indicate increased movement in mutants. Opacity of bars reflects statistical significance ($p < 0.005$). Residue 551 is marked by a vertical dashed line. CFTR domains are shaded and labelled above the plot. Abbreviations: LM, lasso motif; MSD, membrane-spanning domain; NBD, nucleotide-binding domain.

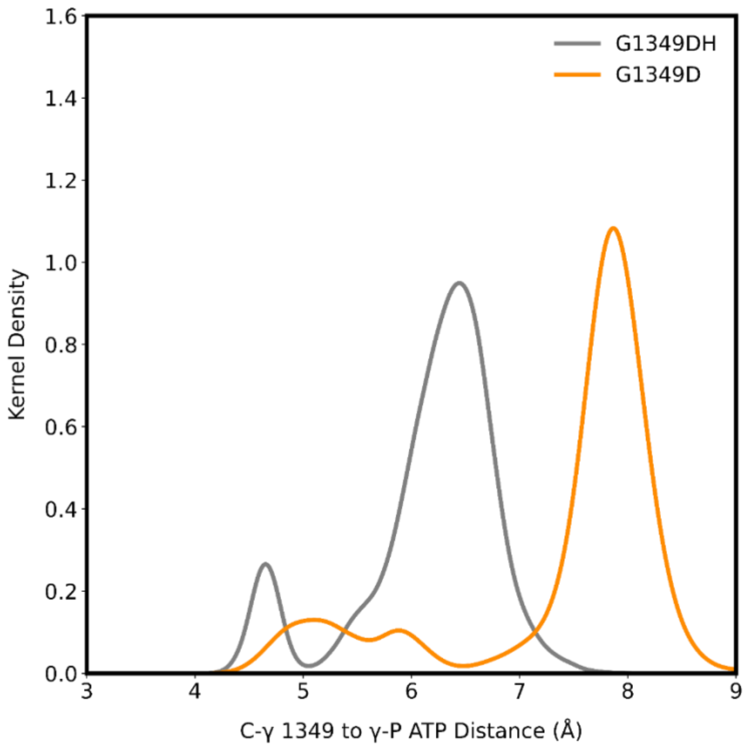


**Figure S4. Aspartate-to-ATP Distances in G1349D CFTR Simulations**

Kernel-density estimates (pooled from five independent 500 ns replicates per system) of the distance (Å) between the C-γ atom of D1349 and the γ-phosphate of ATP in the first ATP-binding site for G1349DH (protonated; grey) and G1349D (deprotonated; orange).

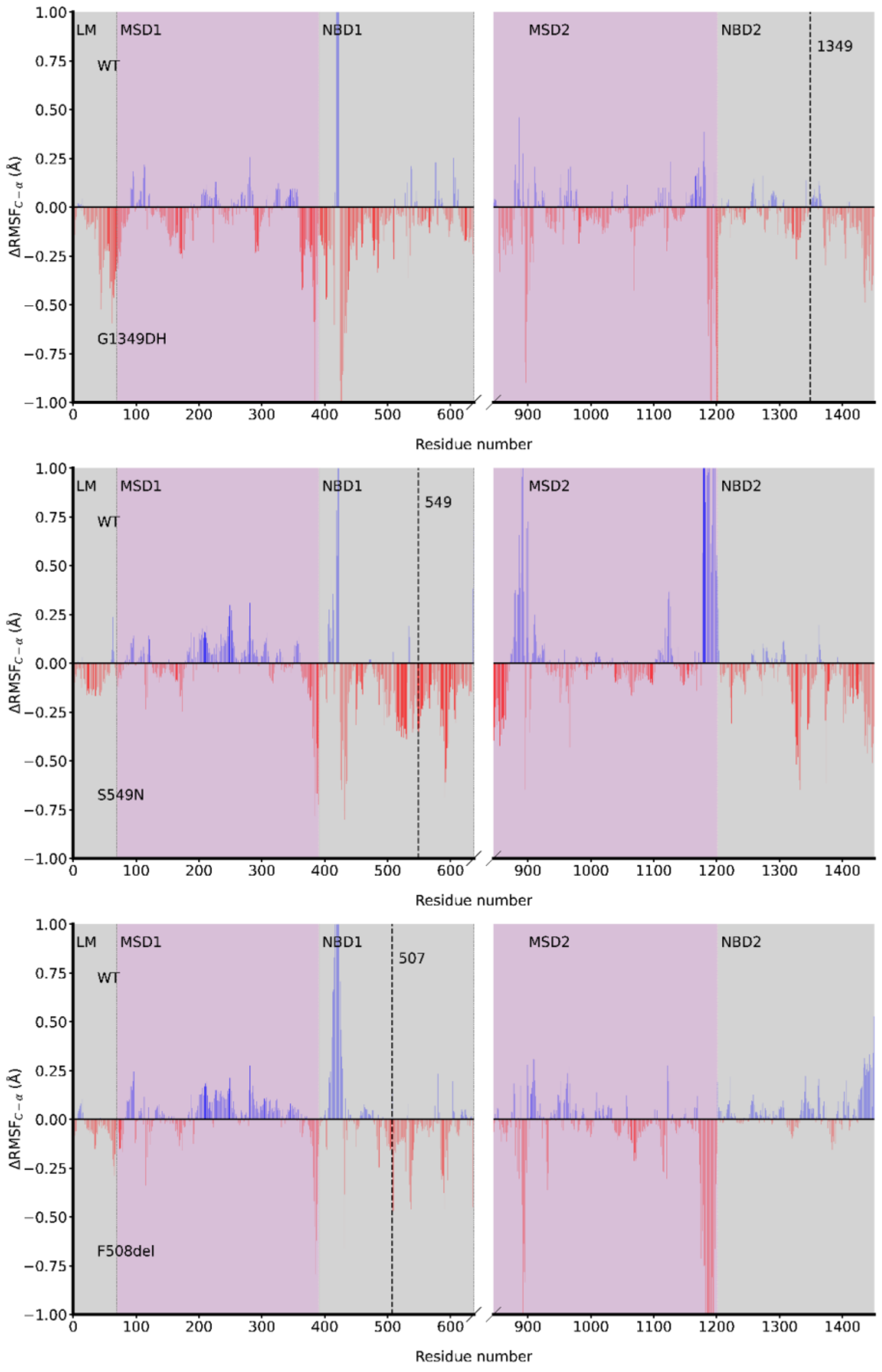


**Figure S5. ΔRMSF analysis between WT and mutant CFTR systems.**

Differences in root-mean-square fluctuation (ΔRMSF = $RMSF_{mutant}$ – $RMSF_{WT}$) for C-α atoms comparing WT CFTR with mutants G1349DH (top panel), S549N (middle panel) and F508del (bottom panel). Blue bars represent regions with greater flexibility in WT, while red bars indicate increased flexibility in the mutants. Opacity of bars reflects statistical significance ($p$ < 0.005). The vertical dashed line marks the positions of the mutations: residue 1349 for G1349D, residue 549 for S549N, and residue 509 for F508del. CFTR domains are shaded and labelled above the plot. Abbreviations: LM, lasso motif; MSD, membrane-spanning domain; NBD, nucleotide-binding domain.

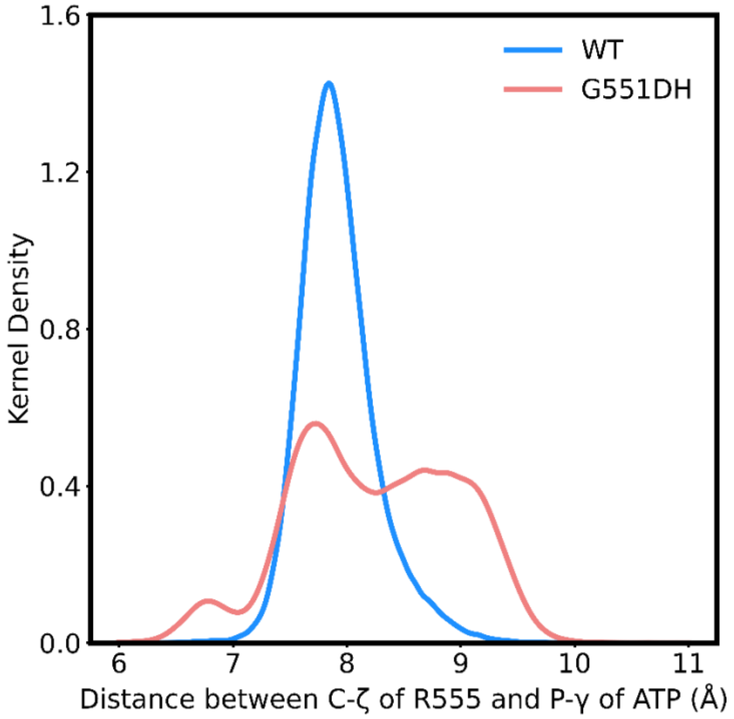


**Figure S6. Residue R555 to ATP Distances in WT and G551DH simulations.**

Kernel-density estimates (pooled from five independent 500 ns replicates) of the distance (Å) between the C-ζ atom of R555 and the γ-phosphate of ATP in ATP-binding site 2 for WT (blue) and G551DH (pink).

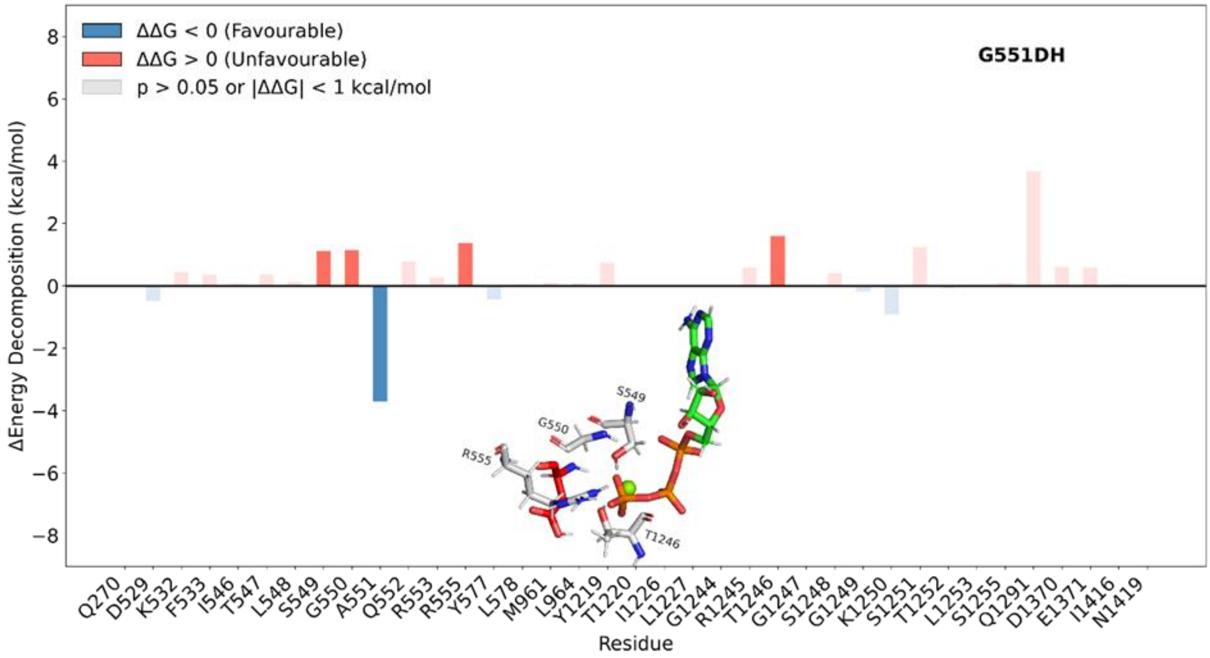


**Figure S7. Per-residue energy decomposition of ATP + $Mg^{2+}$ binding at site 2 in WT versus G551DH.**

Bar plot of per-residue energy differences (ΔΔG = ΔGG551DH – ΔGWT; kcal/mol), where positive values (light orange) indicate destabilising contributions in G551DH (less favourable binding), and negative values (blue) denote stabilising contributions (more favourable binding). Opacity reflects statistical significance across five 500 ns replicates (solid bars indicate p ≤ 0.05, transparent bars *p* > 0.05). Bars for residues with |ΔΔG| < 1 kcal/ mol are shown at reduced opacity to de-emphasise minimal effects. The inset shows the spatial organisation of residues with |ΔΔG| ≥ 1 kcal/mol, rendered as sticks with red carbons for residue 551, and grey carbons for other destabilising residues, green carbons for ATP, and Mg2+ as a green sphere.

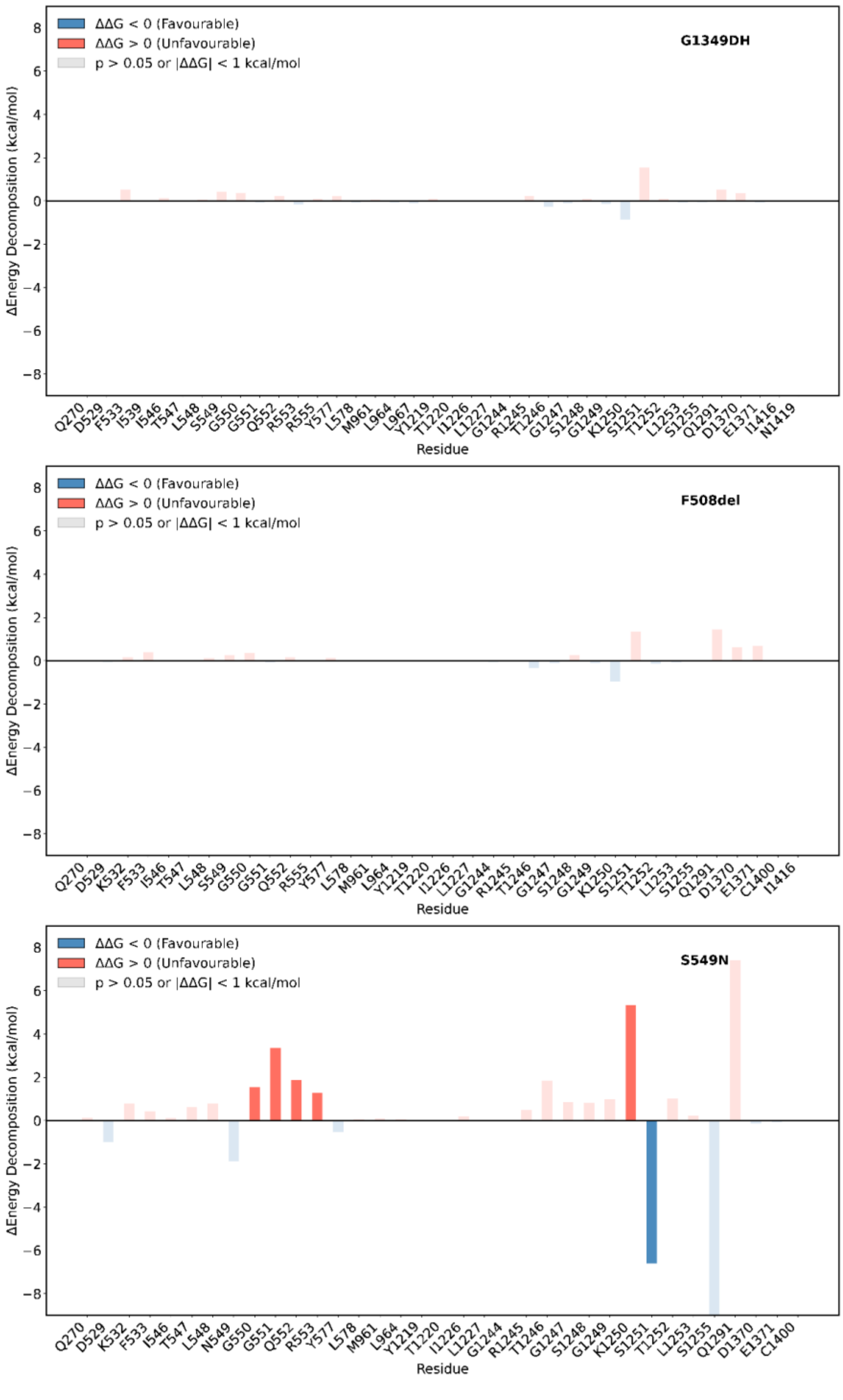


**Figure S8. Per-residue energy decomposition of ATP + $Mg^{2+}$ binding at site 2 in WT versus mutants G1349DH, F508del and S549N.**

Bar plots (top: G1349DH; middle: F508del; bottom: S549N) of per-residue energy differences ($\Delta\Delta G = \Delta G_{mutant} - \Delta G_{WT}$; kcal/mol), where positive values (light orange) indicate stabilising contributions in the mutant (more favourable binding), and negative values (blue) denote destabilising contributions (less favourable binding). Opacity reflects statistical significance across five replicates of 500 ns each (solid bars $p \leq 0.05$; transparent bars $p > 0.05$). Bars for residues with $|\Delta\Delta G| < 1$ kcal/ mol are shown at reduced opacity to de-emphasise minimal effects.

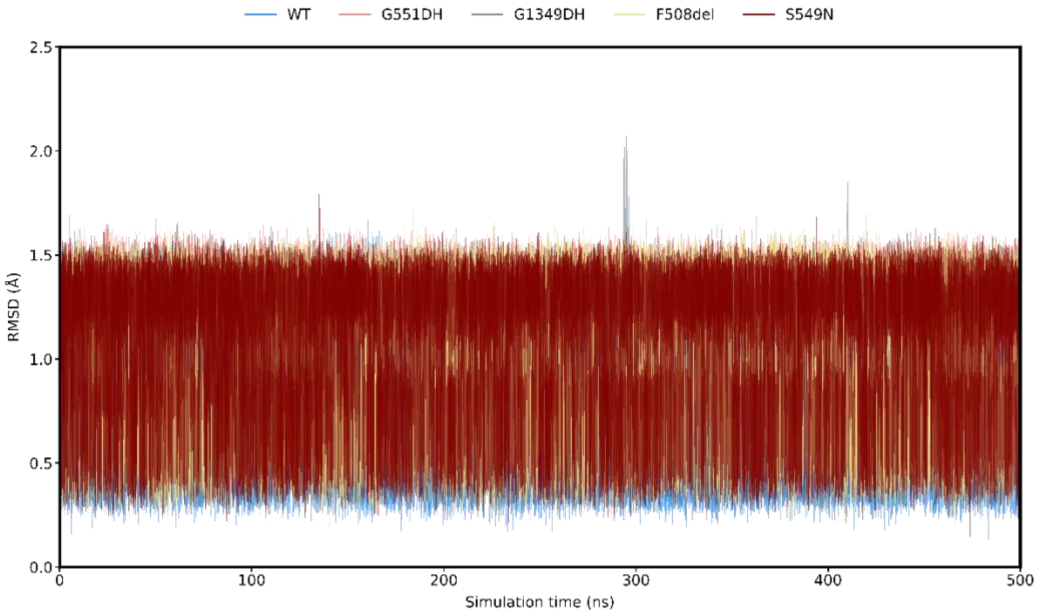


**Figure S9. Time evolution of ivacaftor RMSD in WT and mutant CFTR systems.**

Time traces of ivacaftor RMSD (all ligand atoms) over 0-500 ns MD simulation, pooled from five independent replicates per system. RMSD values (Å) were calculated after aligning each trajectory to the ivacaftor ligand in the first frame. Colours denote WT (blue), G551DH (red), G1349DH-IVA (grey), and F508del (khaki), and S549N (maroon).

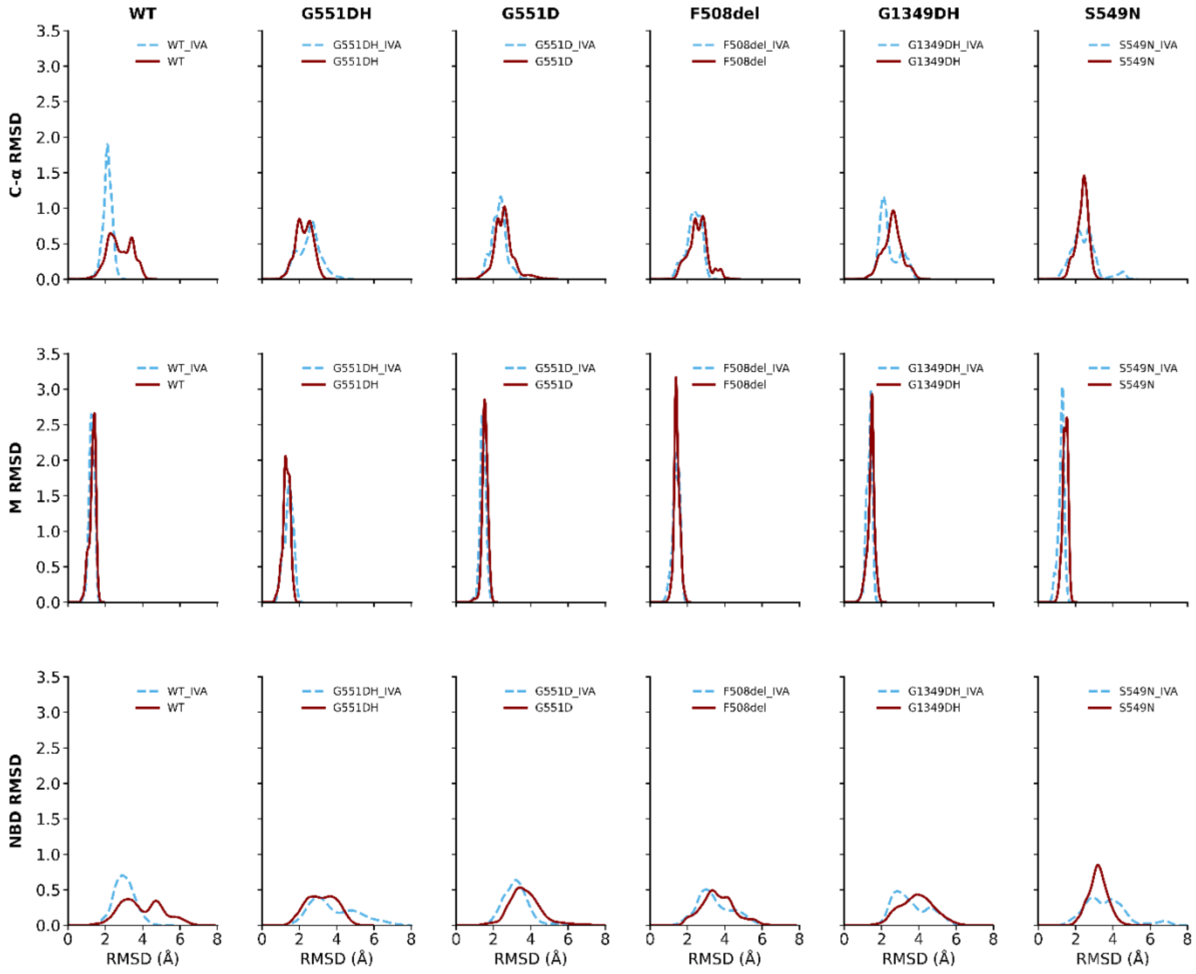


**Figure S10. Effect of ivacaftor on RMSD in WT and mutant CFTR systems.**

Kernel-density estimates of C-α root-mean-square deviation (RMSD) pooled from five independent 500 ns replicates, comparing ivacaftor-bound (blue dashed) and ivacaftor-free (dark red) CFTR systems. The top row shows RMSD for all C-α atoms; the middle row shows the C-α RMSD of transmembrane-helix residues (M1-M12; residues 81-110, 118-150, 177-218, 236-270, 292-392, 330-377, 860-880, 909-959, 966-1012, 1030-1063, 1081-1124, 1125-1162 , as defined in Hwang *et al.*,[55] except M8 (909-959) and M9 (966-1012) as specified here; the bottom row, C-α of the two nucleotide-binding domains (NBD1: residues 391-637; NBD2: residues 1175-1452), each aligned to the transmembrane helices. Individual RMSD traces for all replicate simulations of each system are shown.

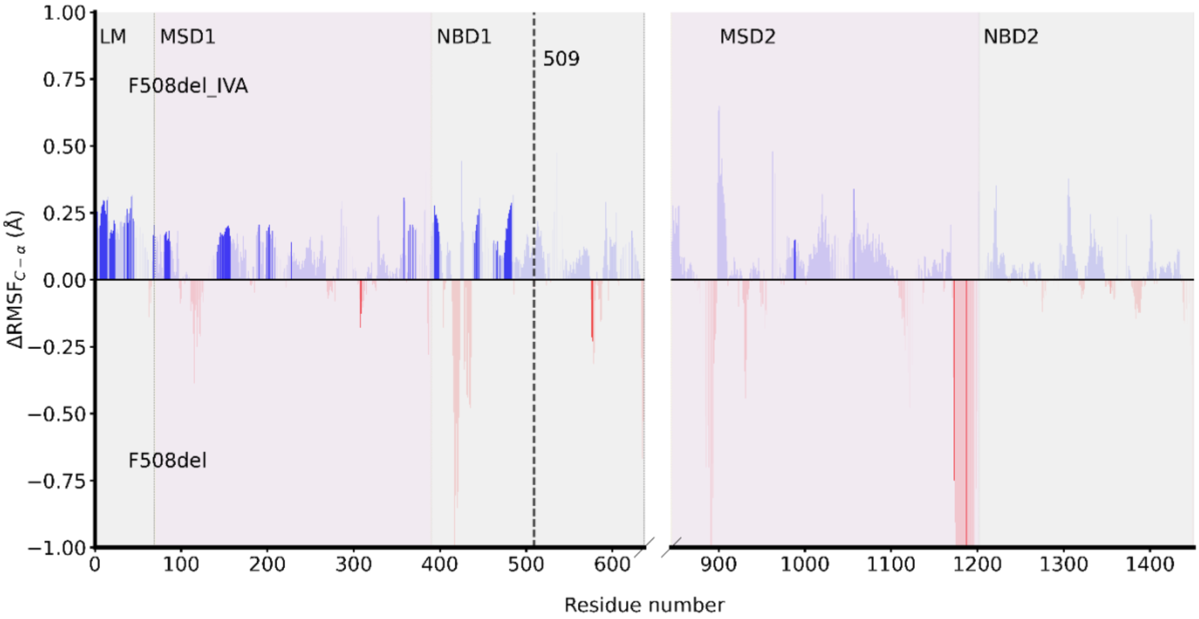


**Figure S11. ΔRMSF analysis of ivacaftor bound and unbound F508del.**

Differences in root-mean-square fluctuation (ΔRMSF = $RMSF_{F508del\_IVA}$ – $RMSF_{F508del}$) for C-α atoms comparing F508del bound by ivacaftor (F508del_IVA) with ivacaftor-free F508del. Blue bars represent regions with greater flexibility in F508del_IVA, while red bars indicate increased movement in F508del. Colour opacity reflects statistical significance ($p < 0.005$). The vertical dashed line marks residue 509 (highlighting the region corresponding to the 508 deletion in this mutant. CFTR structural domains (LM, lasso motif; MSD, membrane-spanning domain; NBD, nucleotide-binding domain) are shaded and labelled above the plot for clarity.

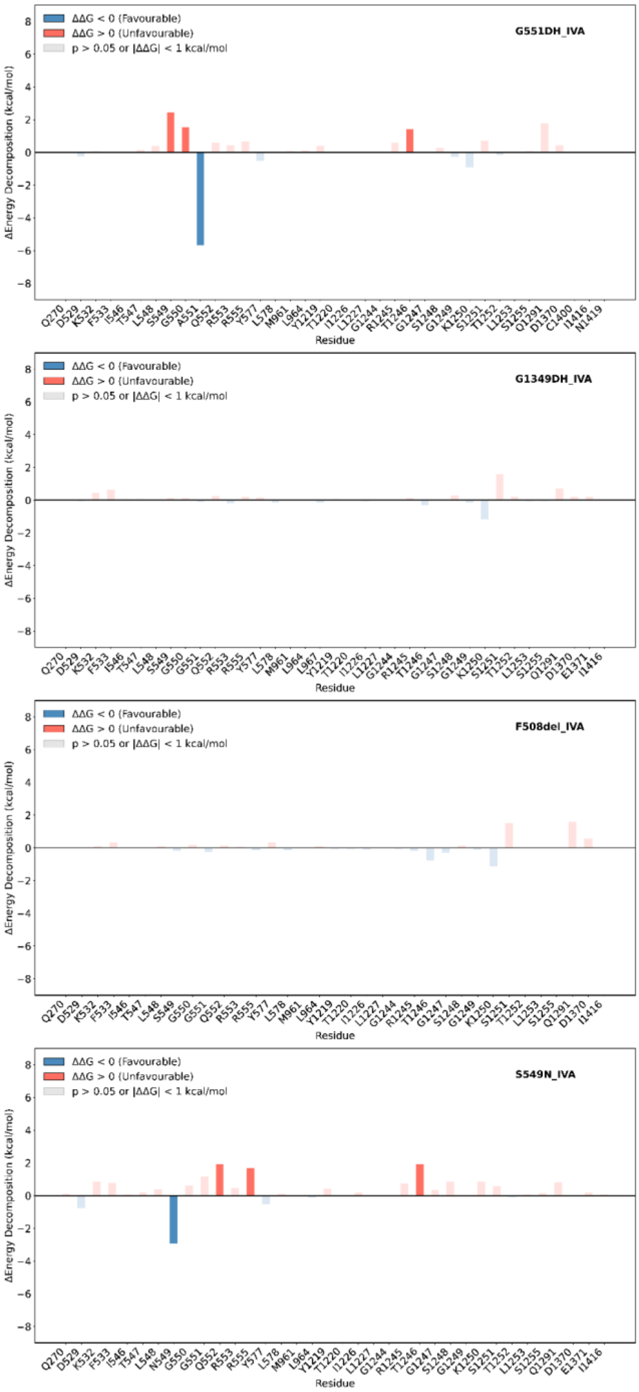


**Figure S12. Per-residue energy decomposition of ATP + $Mg^{2+}$ binding at site 2 in WT versus each ivacaftor-bound CFTR mutant (G551DH_IVA, G1349DH_IVA, S549N_IVA and F508del_IVA).**

Bar plots (top: G551DH_IVA; second: G1349DH_IVA; third: F508del_IVA; bottom: S549N_IVA) of per-residue energy differences ($\Delta\Delta G = \Delta G_{mutant\_IVA} - \Delta G_{WT}$; kcal/mol), where positive values (light orange) indicate stabilising contributions in the mutant (more favourable binding), and negative values (blue) denote destabilising contributions (less favourable binding). Opacity reflects statistical significance across five independent 500 ns replicates per system (solid bars $p \leq 0.05$; transparent bars $p > 0.05$). Bars for residues with $|\Delta\Delta G| < 1$ kcal/mol are shown at reduced opacity to de-emphasise minimal effects.

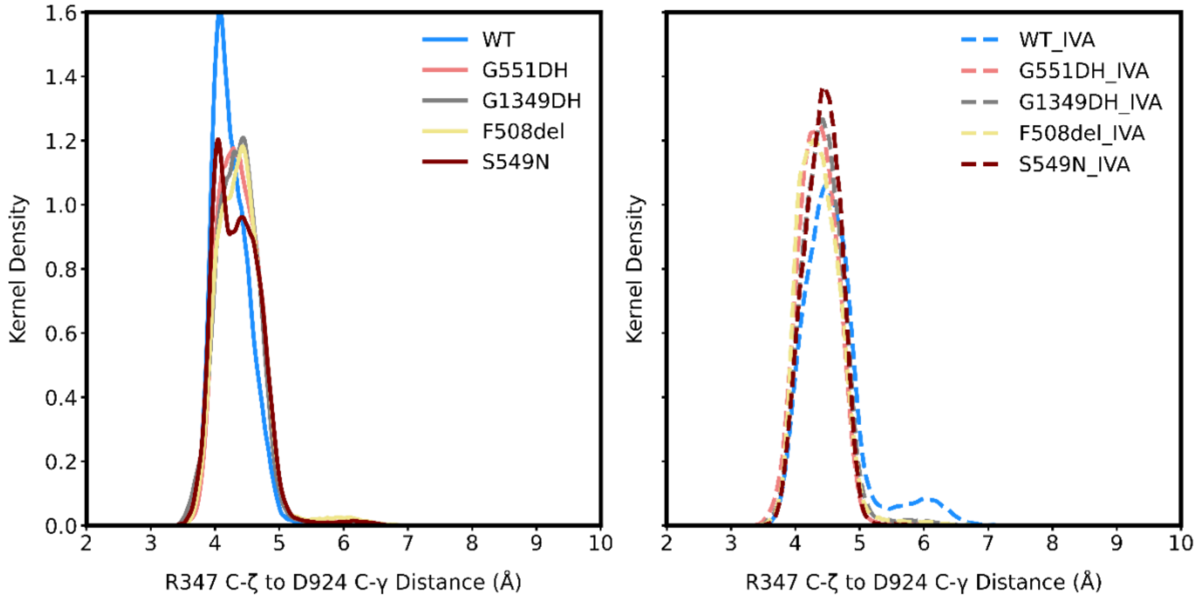


**Figure S13. Pore-domain salt bridge R347-D924 formation in CFTR variants, quantified by distance distributions.**

Kernel-density estimates (pooled from five independent 500 ns replicates per system) of the distance between the C-ζ atom of R347 and the C-γ atom of D924, for simulations without ivacaftor (left) and with ivacaftor (right).

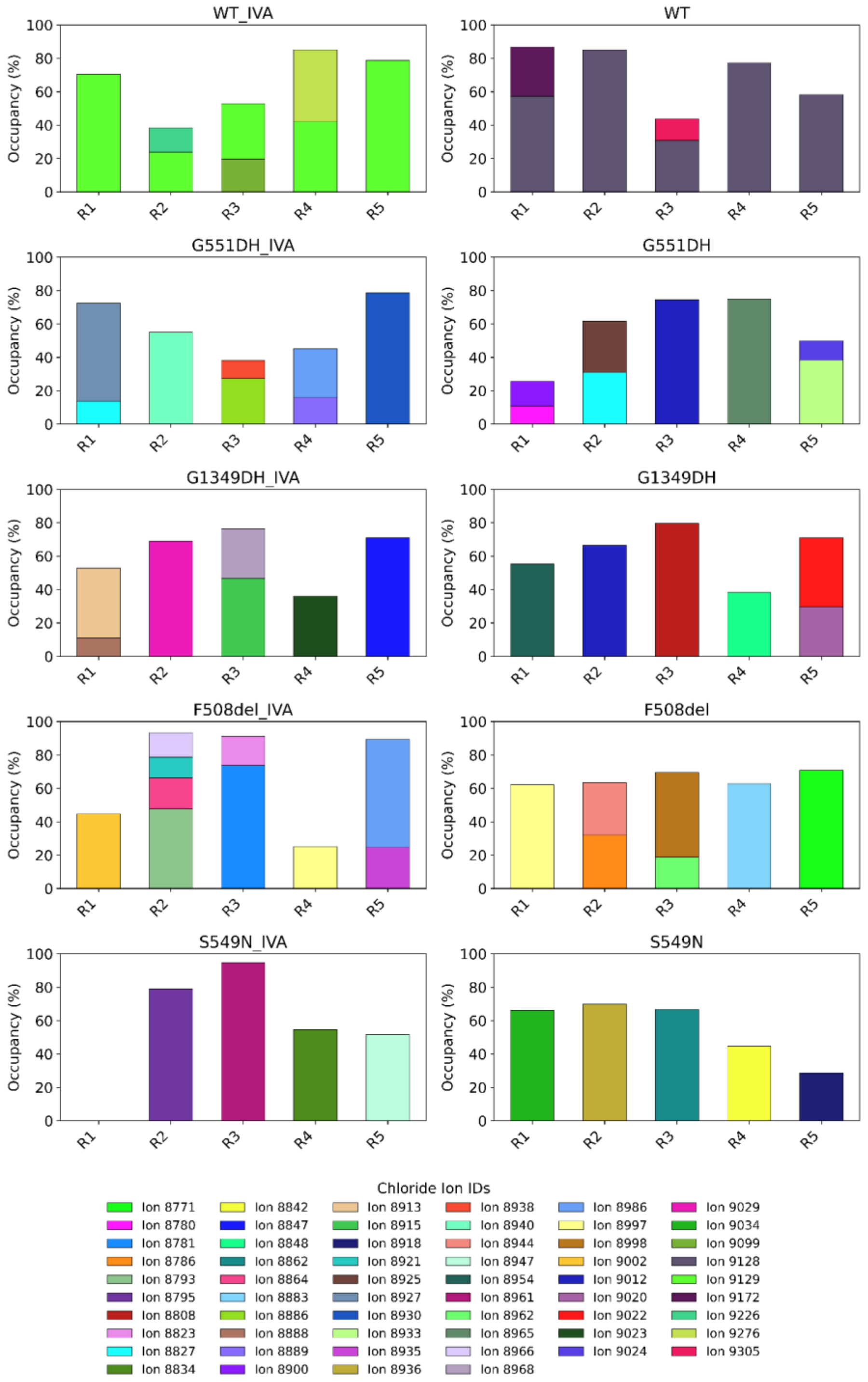


**Figure S14. Chloride ion occupancy across ivacaftor-bound and ivacaftor-free systems.**

Stacked bar plots depict the percentage of simulation frames in which specific chloride ions occupy the pore, grouped by system and replicate (R1-R5). The bars within each subplot are stacked to show how multiple ions may contribute to pore occupancy in a single replicate. Only ions appearing for at least 10% of the total 500 ns trajectory are included. Each colour corresponds to a unique ion ID (see global legend at the bottom).

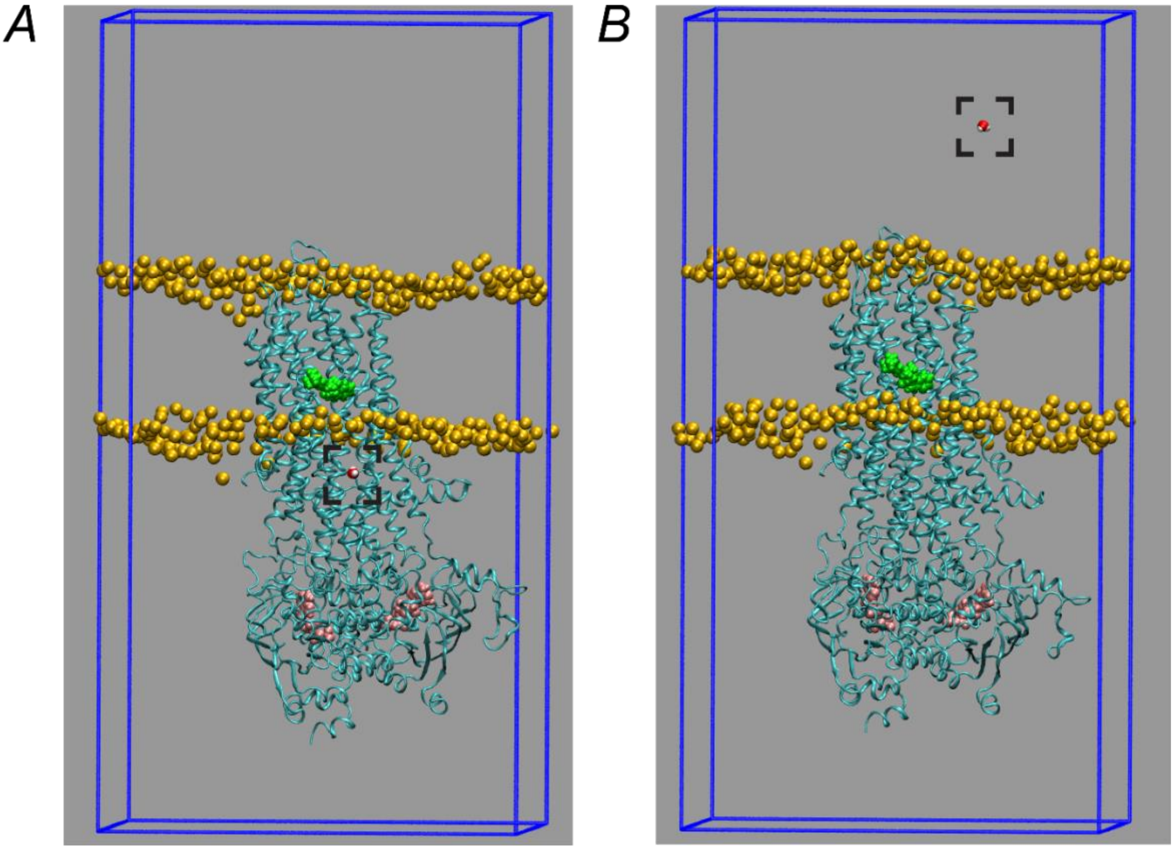


**Figure S15. Water molecule translocation in ivacaftor-bound WT CFTR.**

Representative MD simulation snapshots illustrating a water molecule (marked by a rectangle) moving through the CFTR channel from the intracellular (*A*) to the extracellular side (*B*). The CFTR protein is rendered as a teal cartoon, ivacaftor as green spheres, membrane boundaries as yellow spheres, and ATP-$Mg^{2+}$ as pink spheres. The video file of this movement is available for download at https://github.com/dianaveselu/CFTR/blob/main/bidirectional_water_passage.avi.

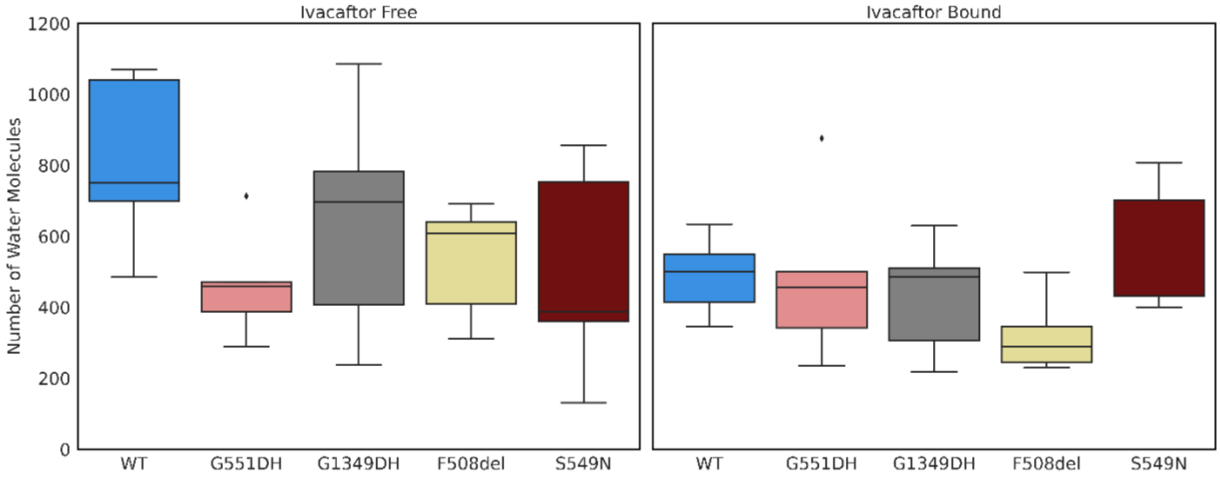


**Figure S16. Water molecules passage through the pore.**

Box plots comparing the number of water molecules passing through the membrane in different CFTR variants under ivacaftor-free (left) and ivacaftor-bound (right) conditions. Each box spans the interquartile range (IQR; $25^{th}$-$75^{th}$ percentile), the horizontal line indicates the median, whiskers extend to the most extreme data points within 1.5 × IQR and dots beyond the whiskers represent outliers.

**Table S1. Binding energies of ATP + $Mg^{2+}$ at ATP-binding site 2 obtained using MM/GBSA.**

| System | IVA-free ΔG ± SD[a] (kcal/mol) | IVA-bound ΔG ± SD[a] (kcal/mol) | p (free vs bound)[b] | p mutant (free) vs WT (free)[b] | p mutant (free) vs WT (bound)[b] |
|---|---|---|---|---|---|
| **WT** | −119.49 ± 9.33 | −109.62 ± 4.75 | 0.08 | — | — |
| **S549N** | −126.27 ± 15.65 | −98.35 ± 7.76 | 0.0122[c] | 0.4351 | 0.0749 |
| **G551DH** | −101.20 ± 9.02 | −110.88 ± 7.38 | 0.1016 | 0.0136 | 0.1137 |
| **F508del** | −109.49 ± 4.91 | −111.16 ± 3.14 | 0.5429 | 0.1182 | 0.5652 |
| **G1349DH** | −112.71 ± 2.99 | −112.15 ± 5.47 | 0.8461 | 0.1848 | 0.259 |

a. Standard deviations are obtained from MM/GBSA calculations on each of the five independent 500 ns replicas for each variant ('IVA-free' - without ivacaftor bound; 'IVA-bound' with ivacaftor bound).

b. p-values obtained from unpaired t-tests comparing (i) ivacaftor-free vs. ivacaftor-bound within each variant, (ii) each ivacaftor-free variant vs ivacaftor-free WT and (iii) ivacaftor-free vs WT ivacaftor-bound.

c. The difference in predicted ATP binding affinity with and without ivacaftor is significant. Residue-level decomposition indicates that two residues destabilise ATP binding in the ivacaftor-bound state (ΔΔG > 1 kcal/mol), with only S1251 significantly so ($p \le 0.05$). It is unclear, however, if and how these changes are related to ivacaftor binding.

**Tabel S2. Cluster occupancy from RMSD-based k-means clustering[a] of ivacaftor after alignment on transmembrane helices.**

| System | Cluster 1 | Cluster 2 | Cluster 3 |
|---|---|---|---|
| **WT_IVA** | 0.5766 | 0.4234 | 0 |
| **G551DH_IVA** | 0.5709 | 0.4291 | 0 |
| **F508del_IVA** | 0.5599 | 0.4399 | 0.0002 |
| **G1349DH_IVA** | 0.5599 | 0.4389 | 0.0013 |
| **S549N_IVA** | 0.5786 | 0.4214 | 0.0001 |

a. RMSD-based k-means clustering ($k = 3$) was performed on ivacaftor coordinates after alignment of trajectories on the C-α atoms of the transmembrane helices (residues 81-110, 118-150, 177-218, 236-270, 292-329, 330-377, 860-880, 909-959, 966-1012, 1030-1063, 1081-1124, 1125-1162). For each system, the table reports the fraction of simulation frames assigned to each cluster, pooled across five independent 500 ns replicas.
